# Detailed profiles of histone modification in male germ line cells of the young and aged mice

**DOI:** 10.1101/635961

**Authors:** Misako Tatehana, Ryuichi Kimura, Kentaro Mochizuki, Noriko Osumi

**Affiliations:** Department of Developmental Neuroscience, Center for Advanced Research and Translational Medicine (ART), Tohoku University School of Medicine, Sendai, Japan; Department of Medical Genetics, Life Sciences Institute, The University of British Columbia, Vancouver, BC V6T 1Z3, Canada

**Keywords:** spermatogenesis, aging, histone modifications

## Abstract

Human epidemiological studies have shown paternal aging as one of the risks for neurodevelopmental disorders such as autism in offspring. A recent study has suggested that factors other than *de novo* mutations due to aging can influence biology of offspring. Here we are focusing on epigenetic alterations in sperm that can influence offspring developmental programs. In this study, we qualitatively and semi-quantitatively evaluated histone modification patterns in male germ line cells throughout spermatogenesis based on immunostaining of testes taken from young (3 months) and aged (12 months) old mice. Although localization patterns were not obviously changed between young and aged testes, some histone modification showed differences in their intensity. Among histone modifications that repress gene expression, H3K9me3 was decreased in the male germ line cells in the aged testis, while H3K27me2/3 was increased. The intensity of H3K27ac, an active mark, was relatively low in the aged testis. Interestingly, H3K27ac was detected in putative sex chromosomes of round spermatids, while other chromosomes were occupied by a repressive mark H3K27me3. Among other histone modifications that activate gene expression, H3K4me2 was drastically decreased in the male germ line cells in the aged testis. H3K79me3 was contrastingly increased and accumulated on the sex chromosomes at M-phase spermatocytes. Therefore, aging induced alterations in the amount of histone modifications, of which patterns were different in individual histone modifications. Moreover, histone modification seems to be differentially regulated by aging on the sex chromosomes and on others. These findings would help elucidate epigenetic mechanisms underlying influence of paternal aging on offspring’s development.

## Introduction

Autism spectrum disorder (ASD) is one of the neurodevelopmental disorders and has been reported to show a pandemic rise in these decades. The most recent survey in the United States shows that one in 59 children aged 8 years have diagnosed as ASD (1). The reason for the rise might include changes in diagnosis, although paternal aging has been suggested for one of the biological factors. For example, meta-analyses with 5,766,794 children born during 1985–2004 in five countries have revealed significant association of advanced paternal age with a risk of ASD in children (2).

Aging induces mutations in the genome of sperm and most of the *de novo* mutations in autistic children are reported to be inherited from fathers (3). However, a study using mathematical model has demonstrated that the risk of psychiatric illness from *de novo* mutations derived from advanced paternal age has been estimated as 10-20% (4). Therefore, some factors other than genetical mutations can influence health and disease of offspring.

One possible mechanism for paternal aging effects on offspring is epigenetic information encoded in sperm. Previously, we have reported that sperm derived from aged mice have more hypomethylated regions in their genome and it possibly affects offspring’s behavior (5). On the other hand, the age-related alteration of histone modification, another major epigenetic factor, remains unclear. Sperm is produced through dynamic and complicated processes, termed spermatogenesis, including meiosis of spermatocytes and morphological changes into mature sperm cells. Considering a frequent chromatin remodeling during spermatogenesis, alteration of histone modification would have a high impact on the spermatogenic processes. Besides, most of the histone protein in the sperm cell nucleus is replaced with protamine, although some remain on genome regions that are functioning in early developmental stages (6). Therefore, features of histone moiety in spermatogenesis would provide essential clues to understand gene regulation after fertilization and subsequent development of offspring. Nevertheless, little known about localization patterns of histone modification in the male germ line cells and their alteration by aging.

Here we first demonstrated comprehensive profiles of major histone modification during murine spermatogenesis, in which individual epigenetic modifications showed dynamic and distinct localization changes. We further examined influence of aging on the histone modifications in each step of spermatogenesis, and found that levels of histone modification were differentially affected by aging during spermatogenesis in the testis. We also noticed accumulation of H3K79me3 on sex chromosomes. Further elucidation of how epigenetic changes due to aging contribute to altered gene regulation in the offspring would provide an insight to understand influence of paternal aging on health and disease of the next generation.

## Materials and methods

### Animals

C57BL/6J mice at 3-months old were purchased from a breeder (Charles River Laboratories, Japan). Some mice were raised up to 5- or 12-months old at Institute for Animal Experimentation Tohoku University Graduate School of Medicine and used for histological and semi-quantitative analyses. All animals were housed in standard cages in a temperature- and humidity-controlled room with a 12-hour light/dark cycle (light on at 8 am), and had free access to standard food and water. All experimental procedures were approved by Ethic Committee for Animal Experiments of Tohoku University Graduate School of Medicine (#2017- MED210), and the animals were treated according to the National Institutes of Health guidance for the care and use of laboratory animals.

### Histological analyses of histone modifications

Histological analysis was performed based on a previous our report (7). Mice were anesthetized with isoflurane and perfused with PBS (pH 7.4) to remove blood. After perfusion, testes were dissected, immersed in 4% PFA in PBS, and cut into half to be post-fixed in 4% PFA overnight at 4°C. After washing in PBS, the testes were cryoprotected by being immersed in 5 % and 10% sucrose in PBS solution for 1 hour each, 20% sucrose in PBS solution overnight, and embedded in OCT compound (Sakura Finetek, Japan). Frozen sections of the testes were cut at 10 μm on a cryostat (Leica, CM3050).

Next, 0.01M citric acid (pH 6.0) solution was heated up using microwave and kept between 90°C and 95°C on a heat plate with gentle stirring. The sections were emerged in the solution for 10 minutes for antigen retrieval. The solution and sections were left to cool down to 40-50°C, and washed in 1×Tris-based saline containing 0.1% Tween20 (TBST) (pH 7.4), and treated with 3% bovine serum albumin and 0.3% Triton X-100 in PBS solution for blocking. The sections were incubated at 4°C overnight with the following primary antibodies at 1/500: i.e., rabbit anti-H3K9me3 (ab8898, abcam), rabbit anti-H3K27me2 (ab24684, abcam), rabbit anti-H3K27me3 (07-449, Millipore), rabbit anti-H3K27ac (ab4729, abcam), rabbit anti-H3K79me2 (ab3594, abcam), rabbit anti-H3K79me3 (ab2621, abcam), mouse anti-SYCP3 (ab97672, abcam). Subsequently, either Cy3-or Alexa488-conjugated secondary antibody (1:500) was applied together with 4’6-diamino-2-phenylindole (DAPI), and the sections were incubated at RT for one hour in a humid chamber. After washing in TBST at room temperature, specimens were mounted in Vectashield (Vector Laboratories, H-1200) or ProLong™ Diamond Antifade Mountant (Invitrogen, P36965). Images were captured using a confocal laser-scanning microscope (LSM780, Carl Zeiss, Jena, Germany).

### Semi-quantitative analyses of histone modifications

Confocal images of the testis sections were semi-quantitatively analyzed by using ImageJ. SYCP3 signals were used to identify stages of spermatogenesis. A mean intensity of DAPI and each histone modification were measured to normalize histone modification signals by DAPI signal.

### DNA Fluorescence in situ hybridization (DNA-FISH) followed by immunofluorescence staining

Antigen retrieval and immunohistochemical staining were performed as described above. Rabbit anti-H3K79me3 (1:500, ab2621, abcam) and Alexa Fluor Plus 647 (1:500, A32728, Invitrogen) were used as primary and secondary antibodies, respectively, and counterstained with DAPI. The sections were mounted in Vectashield. After capturing images using a confocal laser-scanning microscope, cover glasses were removed and the sections were washed in PBS., and immersed in Histo VT One (1:10, 06380-76, Nacalai Tesque) at 90°C for 25 minutes. After washing in PBS, the sections were immersed in 70% and 100% ethanol, dried in the air, and applied with 10 μl probes for X and Y chromosomes (MXY-10, Chromosome Science Lab). The sections were covered with cover slips and heated at 80°C on a heat plate (HDB-1N, AS ONE) for 10 minutes. After 15-hour incubation in a humid chamber at 37°C, the sections were washed in 2×Standard Saline Citrate (SSC) (pH 4.0), and cover slips were removed. The sections were next incubated in 50% formamid-2×SSC at 37°C for 20 minutes and washed in 1×SSC for 15 minutes at room temperature. The sections were mounted in Vectashield. Images were captured using a confocal laser-scanning microscope.

### Statistical analysis

Student’s *t*-test was used to determine statistical significance. Values of *p < 0.05 and **p < 0.01 were considered statistically significant.

## Results

### 1. Morphological profiles showing histone modification patterns during spermatogenesis

Male germ line cells drastically change their morphology during spermatogenesis. Although previous literatures show localization of some histone modifications in these cells (8–10), comprehensive localization patterns are not fully examined. Here, we first investigated localization of major histone modifications during spermatogenesis according to 12 stages based on a previous report (11). This staging helped us to morphologically follow each step of spermatogenesis without any marker staining.

#### (1) Histone modifications that activate gene expression

First, we examined active marks for gene expression, i.e., H3K4me2, H3K4me3, H3K27ac, H3K79me2 and H3K79me3.

##### (i) H3K4me2

H3K4me2 signal was observed in all stages of seminiferous tubules. At stage VIII in which the earliest meiotic spermatocytes (i.e., preleptotene cells) were located near the periphery of the tubule, a signal of H3K4me2 was detected in the whole nucleus of preleptotene cells, though constitutive heterochromatin did not show the signal (Fig. 1a, arrowhead 1). The signal continued throughout meiosis (Fig. 1a, arrowheads 2-6). In the pachytene cells of stage X, a strong signal was detected on the XY body (Fig. 1a, arrowhead 7). The accumulation of H3K4me2 on the XY body disappeared in the M-phase spermatocytes at stage XII, and only weak signal was observed in the chromosomes (Fig. 1a, arrowhead 8). In round spermatids from stage I to VIII and elongated spermatids at stage X, the signal was widely detected in the nucleus, but not in the chromocenter (constitutive heterochromatin) (Fig. 1a, arrowheads 9-12). Thus, localization of H3K4me2 modification seems to start from preleptotene cells, and to accumulate on the XY body in late pachytene cells. Through spermatogenesis, H3K4me2 was not detected in the chromocenter (Fig. 1b).

**Fig. 1.**
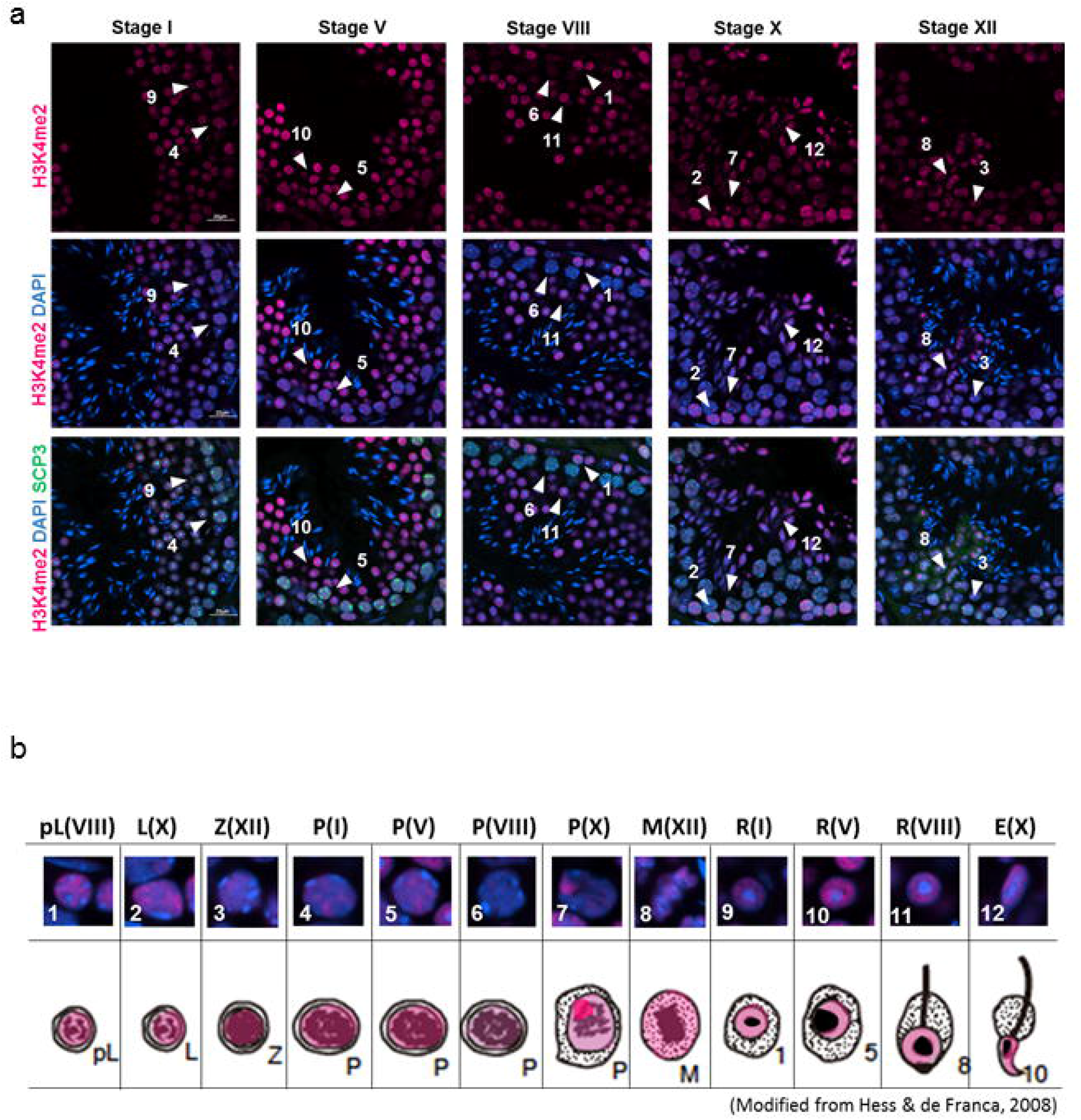
Localization of H3K4me2 in male germ line cells in the young (3 M) testis. (a) Representative confocal images of H3K4me2 (magenta) and SCP3 (green) in the seminiferous epithelium in stages I, V, VIII, X and XII. Nuclei were counterstained with DAPI (blue). The number with arrowhead means as the followings: 1; Preleptotene spermatocyte at stage VIII, 2; Leptotene spermatocyte at stage X, 3; Zygotene spermatocyte at stage XII, 4-7; Pachytene spermatocyte at stage I, V, VIII and X respectively, M; M phase cells at stage XII, 9-11; Round spermatid at stage I, V and VIII respectively, 12; Elongated spermatid at stage X. Scale bar: 20 μm. (b) Magnified images of the cells indicated by arrowheads in (a), and subcellular localization of H3K4me2 in the young testis is shown in magenta in a graphical summary. pL: Preleptotene spermatocyte, L: Leptotene spermatocyte, Z: Zygotene spermatocyte, P: Pachytene spermatocyte, M: M phase, R: Round spermatid, E: Elongated spermatid. Each parenthesis represents the stage of spermatogenesis.

##### (ii) H3K4me3

While H3K4me2 was observed from preleptotene cells (Fig. 1b), H3K4me3 signal was not detected from preleptotene cells to pachytene cells (Fig. 2a, arrowheads 1-7). H3K4me3 was firstly detected in M-phase spermatocytes in stage XII seminiferous tubules (Fig. 2a, arrowhead 8). The signal was broadly distributed on condensed chromosomes (Fig. 2b, arrowhead 8). After meiotic division II, the signal was detected in round spermatids at stage I to VIII, though there was no signal in the chromocenter (Fig. 2a, arrowheads 9-11). In elongated spermatids, the signal was weak and widely detected in whole nucleus (Fig. 2a, arrowhead 12). Thus, localization of H3K4me3 modification appeared later stage of meiosis, and the signal was weak compare to H3K4me2 modification (Fig. 2b).

**Fig. 2.**
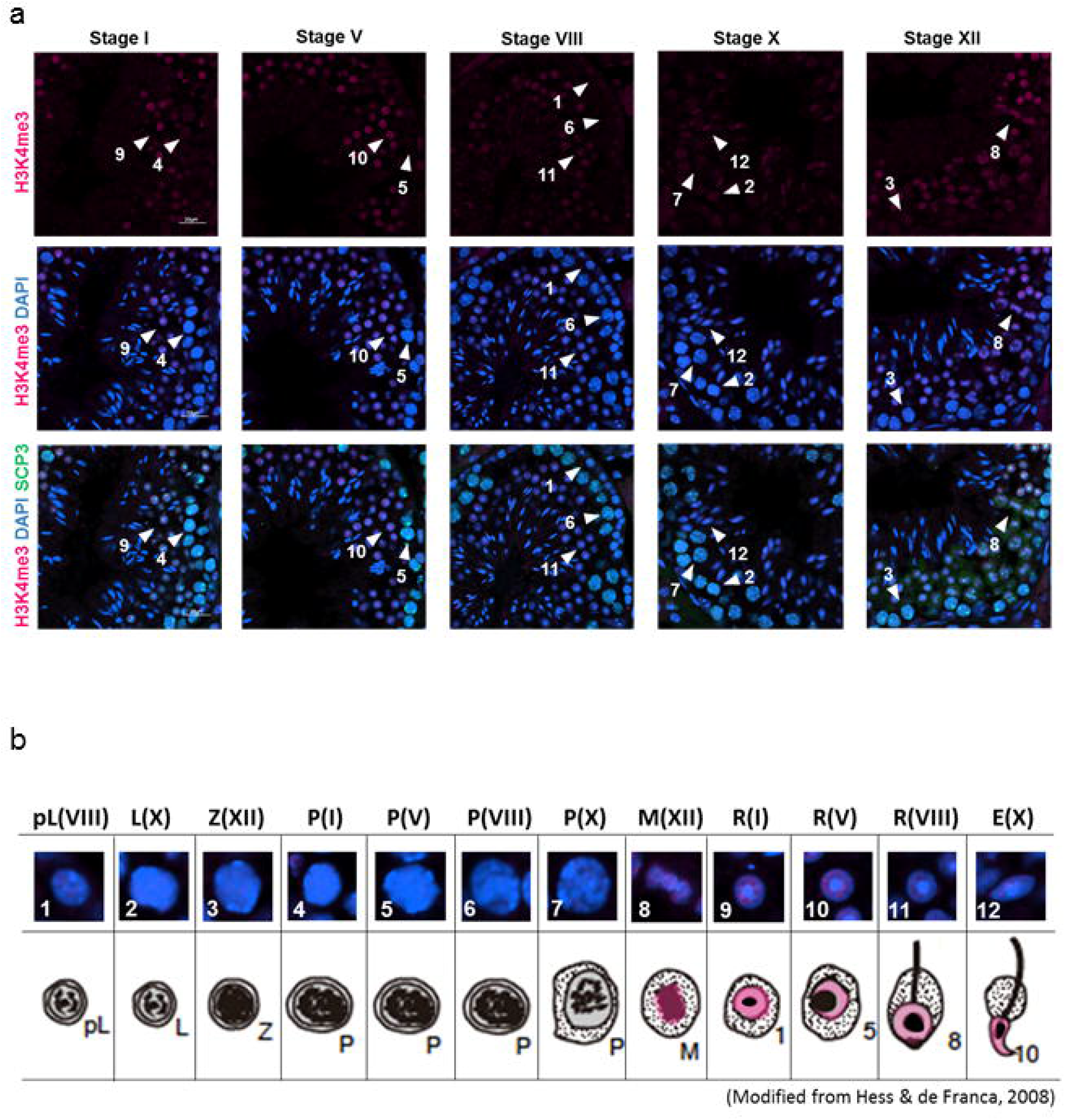
Localization of H3K4me3 in male germ line cells in the young (3 M) testis. (a) Representative confocal images of H3K4me3 (magenta) and SCP3 (green) in the seminiferous epithelium in stages I, V, VIII, X and XII. Nuclei were counterstained with DAPI (blue). The number with arrowhead means as the followings: 1; Preleptotene spermatocyte at stage VIII, 2; Leptotene spermatocyte at stage X, 3; Zygotene spermatocyte at stage XII, 4-7; Pachytene spermatocyte at stage I, V, VIII and X respectively, M; M phase cells at stage XII, 9-11; Round spermatid at stage I, V and VIII respectively, 12; Elongated spermatid at stage X. Scale bar: 20 μm. (b) Magnified images of the cells indicated by arrowheads in (a), and subcellular localization of H3K4me3 in the young testis is shown in magenta in a graphical summary. pL: Preleptotene spermatocyte, L: Leptotene spermatocyte, Z: Zygotene spermatocyte, P: Pachytene spermatocyte, M: M phase, R: Round spermatid, E: Elongated spermatid. Each parenthesis represents the stage of spermatogenesis.

##### (iii) H3K27ac

H3K27ac signal was observed in all stages of seminiferous tubules. At stage VIII in which the earliest meiotic cells (i.e., preleptotene cells) were located near the periphery of the tubule, strong signals of H3K27ac were detected in whole nuclei of preleptotene cells (Fig. 3a, arrowhead 1). While the signal continued throughout meiosis, the intensity dramatically decreased when preleptotene cells differentiated into leptotene cells (Fig. 3a, arrowhead 2-5). In the pachytene cells of stage VIII, a strong signal was detected on the XY body (Fig. 3a, arrowhead 6). The accumulation of H3K27ac on the XY body disappeared in the pachytene cells of stage X and M-phase spermatocytes at stage XII, and only weak signal was observed in the whole nucleus or chromosomes (Fig. 3a, arrowheads 7-8). In round spermatids from stage I to V, the signal was widely detected in the nucleus, especially in the peri-chromocenter (putative sex chromosome) (Fig. 3a, arrowheads 9-10). The strong signal on the peri-chromocenter disappeared in round spermatids of stage VIII (Fig. 3a, arrowhead 11). H3K27ac signal increased again in elongated spermatids, suggesting requirement of H3K27ac for spermiogenesis (Fig. 3a, arrowhead 12). Thus, localization of H3K27ac modification seems to start from preleptotene cells, and to accumulate on the XY body in late pachytene cells and on the putative sex chromosome during early to middle stages of round spermatids (Fig. 3b).

**Fig. 3.**
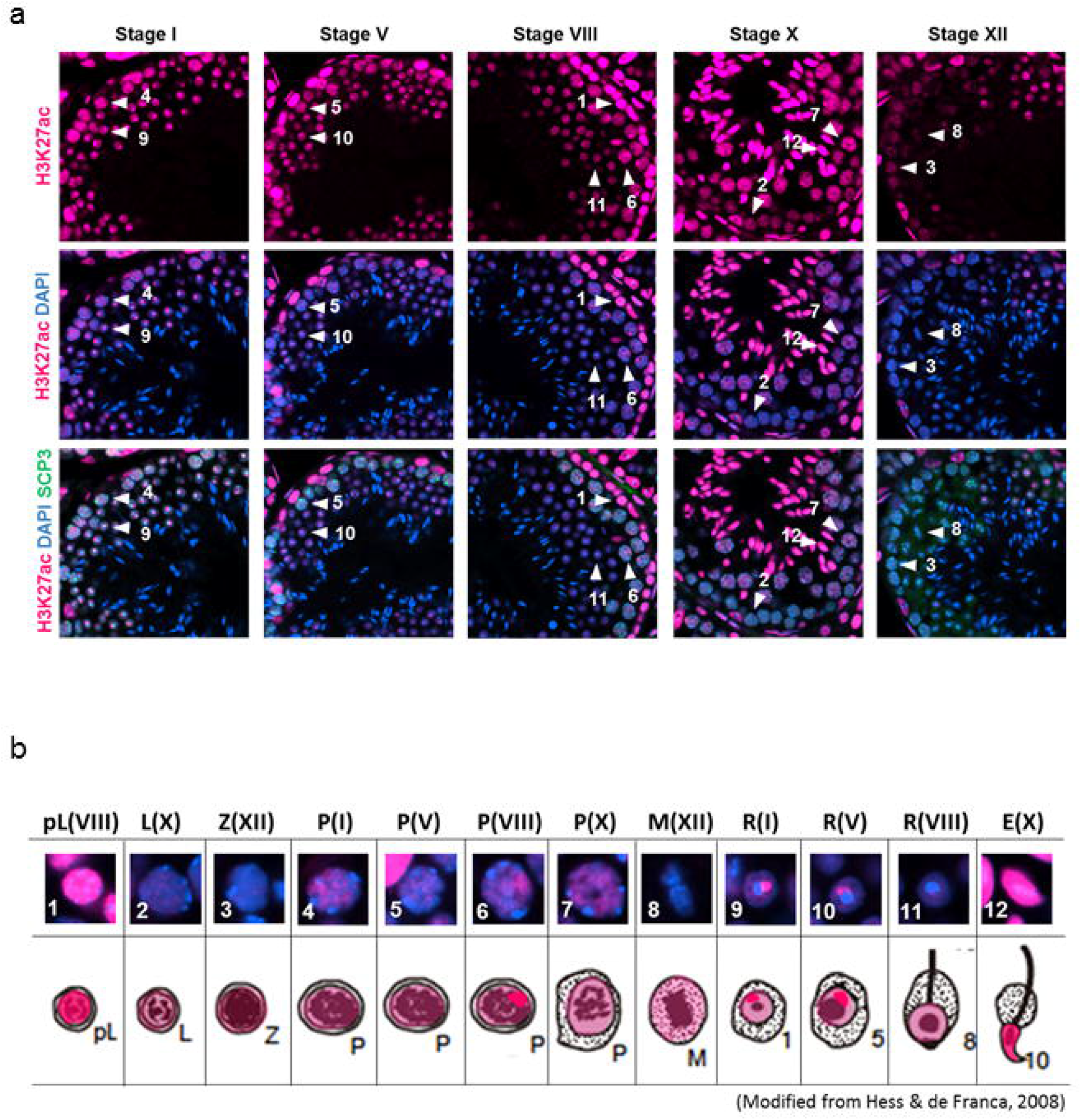
Localization of H3K27ac in male germ line cells in the young (3 M) testis. (a) Representative confocal images of H3K27ac (magenta) and SCP3 (green) in the seminiferous epithelium in stages I, V, VIII, X and XII. Nuclei were counterstained with DAPI (blue). The number with arrowhead means as the followings: 1; Preleptotene spermatocyte at stage VIII, 2; Leptotene spermatocyte at stage X, 3; Zygotene spermatocyte at stage XII, 4-7; Pachytene spermatocyte at stage I, V, VIII and X respectively, M; M phase cells at stage XII, 9-11; Round spermatid at stage I, V and VIII respectively, 12; Elongated spermatid at stage X. Scale bar: 20 μm. (b) Magnified images of the cells indicated by arrowheads in (a), and subcellular localization of H3K27ac in the young testis is shown in magenta in a graphical summary. pL: Preleptotene spermatocyte, L: Leptotene spermatocyte, Z: Zygotene spermatocyte, P: Pachytene spermatocyte, M: M phase, R: Round spermatid, E: Elongated spermatid. Each parenthesis represents the stage of spermatogenesis.

##### (iv) H3K79me2

In pachytene cells from stage VIII to stage X, the signal was widely detected in the nucleus (Fig. 4a, arrowheads 1 and 2). In M-phase spermatocytes at stage XII, the signal was broadly observed on the condensed chromosomes (Fig. 4a, arrowhead 3). In round spermatids at stage I, a weak signal of H3K79me2 was observed in the most of the nucleus except the chromocenter (Fig. 4a, arrowhead 4). Interestingly, a strong signal of H3K79me2 was observed in the peri-chromocenter (a putative sex chromosome) in round spermatids from stage V to VIII (Fig. 4a, arrowheads 5 and 6), while no signal was observed on the chromocenter itself. In elongated spermatids at stage X, the strong signal on the putative sex chromosome disappeared (Fig. 4a, arrowhead 7). Thus, localization of H3K79me2 modification seems to start from the late pachytene cells, and to accumulate on the putative sex chromosome during most stages of round spermatids (Fig. 4b).

**Fig. 4.**
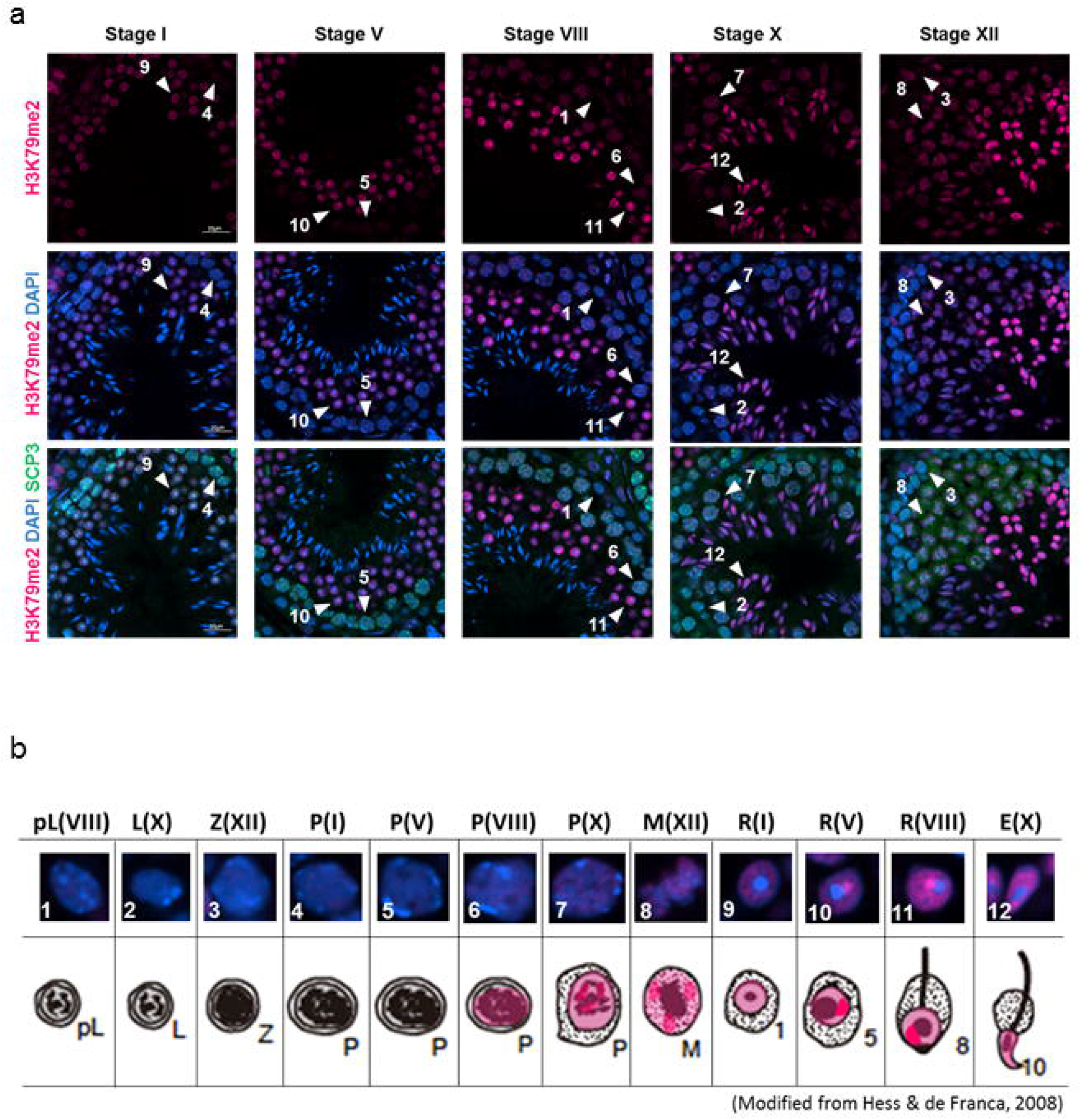
Localization of H3K79me2 in male germ line cells in the young (3 M) testis. (a) Representative confocal images of H3K79me2 (magenta) and SCP3 (green) in the seminiferous epithelium in stages I, V, VIII, X and XII. Nuclei were counterstained with DAPI (blue). The number with arrowhead means as the followings: 1; Preleptotene spermatocyte at stage VIII, 2; Leptotene spermatocyte at stage X, 3; Zygotene spermatocyte at stage XII, 4-7; Pachytene spermatocyte at stage I, V, VIII and X respectively, M; M phase cells at stage XII, 9-11; Round spermatid at stage I, V and VIII respectively, 12; Elongated spermatid at stage X. Scale bar: 20 μm. (b) Magnified images of the cells indicated by arrowheads in (a), and subcellular localization of H3K79me2 in the young testis is shown in magenta in a graphical summary. pL: Preleptotene spermatocyte, L: Leptotene spermatocyte, Z: Zygotene spermatocyte, P: Pachytene spermatocyte, M: M phase, R: Round spermatid, E: Elongated spermatid. Each parenthesis represents the stage of spermatogenesis.

##### (v) H3K79me3

While H3K79me2 was observed from the late pachytene cells (Fig. 4b), H3K79me3 signal was not detected in any pachytene cells. H3K79me3 was first detected in M-phase spermatocytes in stage XII seminiferous tubules (Fig. 5a, arrowhead 8). Unlike H3K79me2, H3K79me3 signal was not distributed on all condensed chromosomes, but on specific chromosomes (Fig. 5a, arrowhead 8). After meiotic division II, the signal was not detected in round spermatids at stage I (Fig. 5a, arrowhead 9). However, in round spermatids from stage V to VIII, H3K79me3 signal was observed again in the nucleus broadly and strong signal was detected in the peri-chromocenter (a putative sex chromosome) (Fig. 5a, arrowheads 10-11). In elongated spermatids at stage X, the signal was widely observed in the nucleus (Fig. 5a, arrowhead 12). Thus, H3K79me3 modification appeared from the late stage of meiosis, and accumulated on the putative sex chromosome of round spermatids. A specific area of condensed chromosomes showed strong signal of H3K79me3 during M-phase (Fig. 5b).

**Fig. 5.**
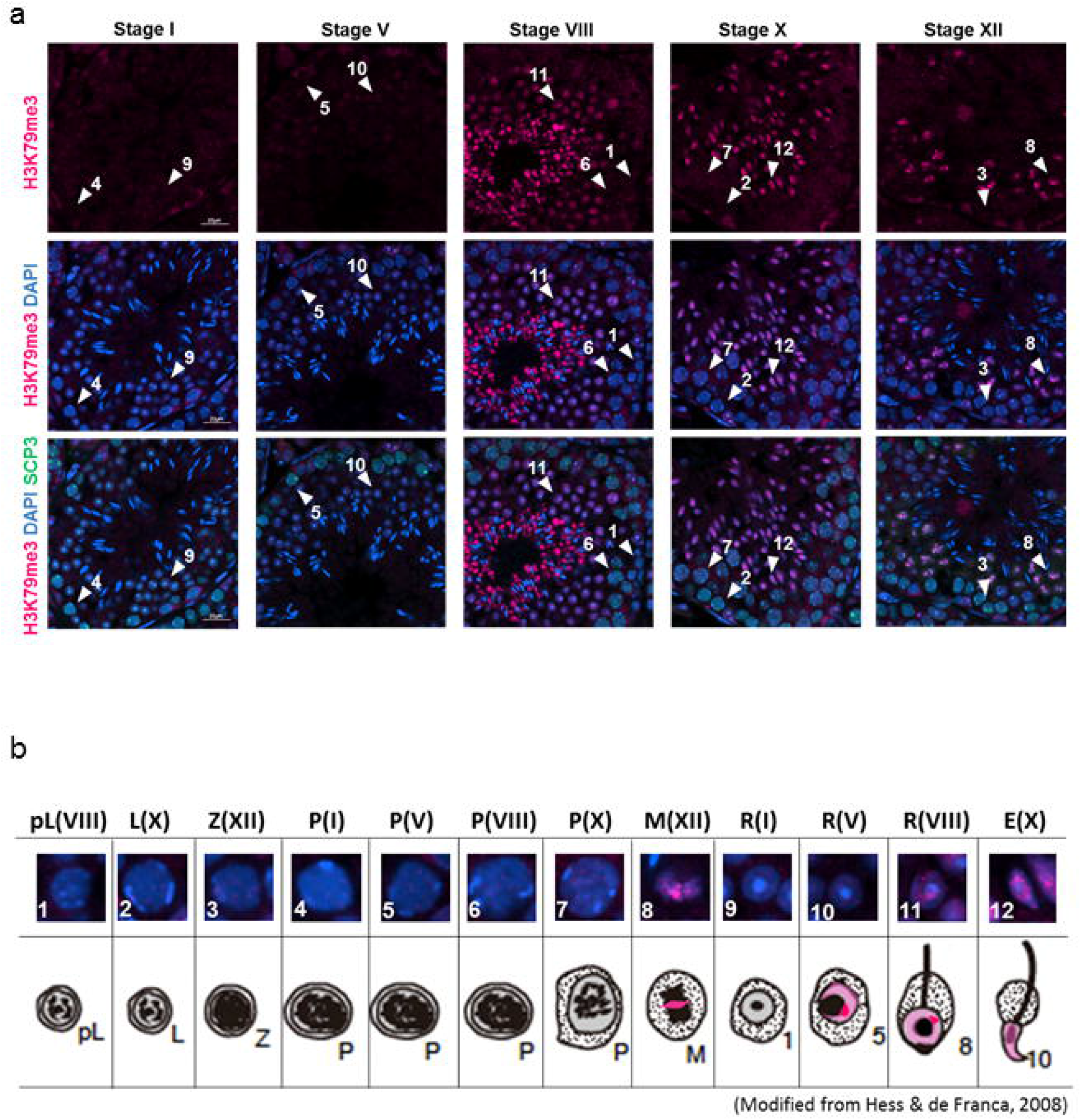
Localization of H3K79me3 in male germ line cells in the young (3 M) testis. (a) Representative confocal images of H3K79me3 (magenta) and SCP3 (green) in the seminiferous epithelium in stages I, V, VIII, X and XII. Nuclei were counterstained with DAPI (blue). The number with arrowhead means as the followings: 1; Preleptotene spermatocyte at stage VIII, 2; Leptotene spermatocyte at stage X, 3; Zygotene spermatocyte at stage XII, 4-7; Pachytene spermatocyte at stage I, V, VIII and X respectively, M; M phase cells at stage XII, 9-11; Round spermatid at stage I, V and VIII respectively, 12; Elongated spermatid at stage X. Scale bar: 20 μm. (b) Magnified images of the cells indicated by arrowheads in (a), and subcellular localization of H3K79me3 in the young testis is shown in magenta in a graphical summary. pL: Preleptotene spermatocyte, L: Leptotene spermatocyte, Z: Zygotene spermatocyte, P: Pachytene spermatocyte, M: M phase, R: Round spermatid, E: Elongated spermatid. Each parenthesis represents the stage of spermatogenesis.

#### (2) Histone modifications that inhibit gene expression

Next, we examined repressive marks, H3K9me3, H3K27me2 and H3K27me3.

##### (i) H3K9me3

H3K9me3 was detected in all cell types examined in this study. Broad nuclear signals were observed from preleptotene cells at stage VIII to pachytene cells at stage V with strong dot signals in the periphery of the nucleus (Fig. 6a, arrowheads 1-5). In pachytene cells at stage VIII and stage X, only dot-like signals were detected (Fig. 6a, arrowheads 6-7). Those dotted signals matched with intense DAPI signals, indicating heterochromatin areas. Consistent with a previous report (12), H3K9me3 exceptionally localized on the sex chromosome of M-phase spermatocytes at stage XII (Fig. 6a, arrowhead 8). In round spermatids at stage I to elongated spermatids at stage X, the signal was detected on the chromocenter and peri-chromocenter (putative sex chromosome) (Fig. 6a, arrowheads 9-12). Thus, localization of H3K9me3 modification seems to start from preleptotene cells, and to accumulate on the sex chromosome in M-phase spermatocytes until late round spermatids (Fig. 6b).

**Fig. 6.**
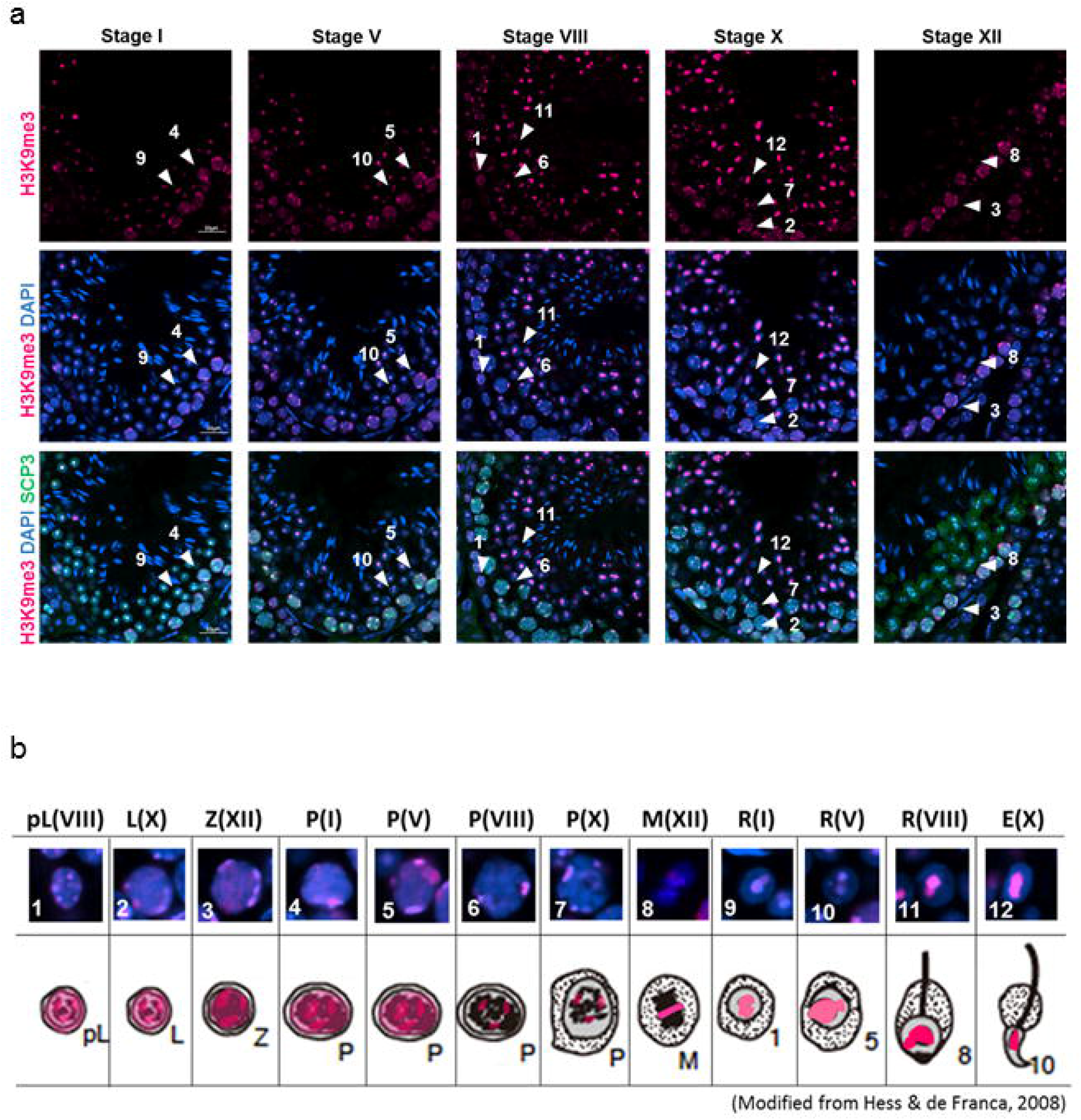
Localization of H3K9me3 in male germ line cells in the young (3 M) testis. (a) Representative confocal images of H3K9me3 (magenta) and SCP3 (green) in the seminiferous epithelium in stages I, V, VIII, X and XII. Nuclei were counterstained with DAPI (blue). The number with arrowhead means as the followings: 1; Preleptotene spermatocyte at stage VIII, 2; Leptotene spermatocyte at stage X, 3; Zygotene spermatocyte at stage XII, 4-7; Pachytene spermatocyte at stage I, V, VIII and X respectively, M; M phase cells at stage XII, 9-11; Round spermatid at stage I, V and VIII respectively, 12; Elongated spermatid at stage X. Scale bar: 20 μm. (b) Magnified images of the cells indicated by arrowheads in (a), and subcellular localization of H3K9me3 in the young testis is shown in magenta in a graphical summary. pL: Preleptotene spermatocyte, L: Leptotene spermatocyte, Z: Zygotene spermatocyte, P: Pachytene spermatocyte, M: M phase, R: Round spermatid, E: Elongated spermatid. Each parenthesis represents the stage of spermatogenesis.

##### (ii) H3K27me2

H3K27me2 was detected in the nucleus of leptotene cells in stage X to pachytene cells in stage X in a salt-and-pepper manner (Fig. 7a, arrowheads 2-7). In M-phase spermatocytes at stage XII, the signal was broadly observed on all chromosomes (Fig. 7a, arrowhead 8). In round spermatids at stage I, strong dot-like signals were observed on the chromocenter in addition to weak and broad signals in the nucleus (Fig. 7a, arrowhead 9). The dot-like signals were observed until in round spermatids of stage VIII (Fig. 7a, arrowheads 10-11), and disappeared in elongated spermatids at stage X (Fig. 7a, arrowhead 12). Thus, localization of H3K27me2 modification stated from leptotene cells, and accumulated on chromocenter of all stages of round spermatids, though the signal was relatively weak through spermatogenesis (Fig. 7b).

**Fig. 7.**
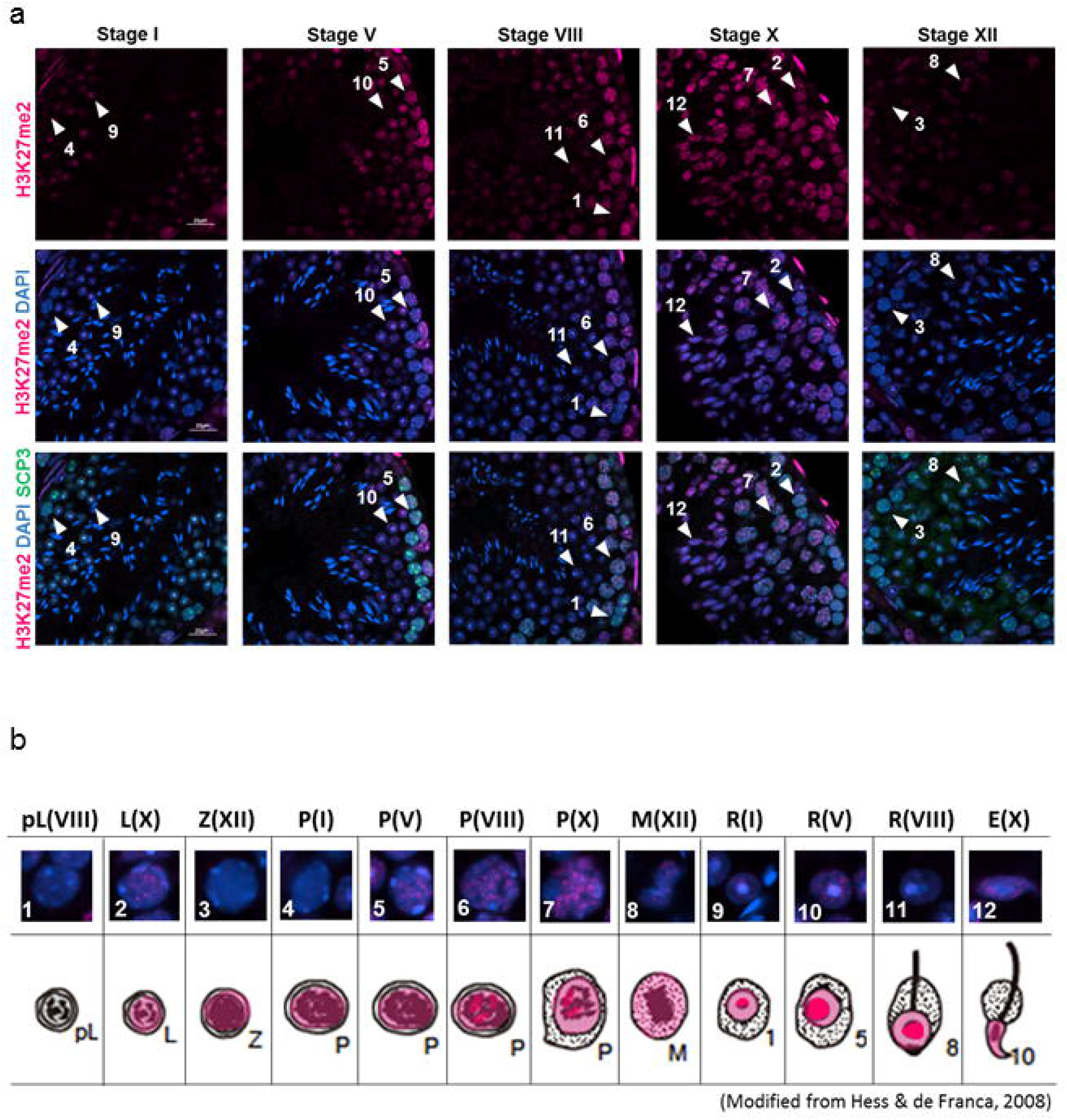
Localization of H3K27me2 in male germ line cells in the young (3 M) testis. (a) Representative confocal images of H3K27me2 (magenta) and SCP3 (green) in the seminiferous epithelium in stages I, V, VIII, X and XII. Nuclei were counterstained with DAPI (blue). The number with arrowhead means as the followings: 1; Preleptotene spermatocyte at stage VIII, 2; Leptotene spermatocyte at stage X, 3; Zygotene spermatocyte at stage XII, 4-7; Pachytene spermatocyte at stage I, V, VIII and X respectively, M; M phase cells at stage XII, 9-11; Round spermatid at stage I, V and VIII respectively, 12; Elongated spermatid at stage X. Scale bar: 20 μm. (b) Magnified images of the cells indicated by arrowheads in (a), and subcellular localization of H3K27me2 in the young testis is shown in magenta in a graphical summary. pL: Preleptotene spermatocyte, L: Leptotene spermatocyte, Z: Zygotene spermatocyte, P: Pachytene spermatocyte, M: M phase, R: Round spermatid, E: Elongated spermatid. Each parenthesis represents the stage of spermatogenesis.

##### (iii) H3K27me3

H3K27me3 was detected in all cell types examined in this study except preleptotene cells. In leptotene cells at stage X to pachytene cells in stage X, the signal was broadly and slightly detected in the nucleus (Fig. 8a, arrowheads 2-7). In M-phase spermatocytes at stage XII, the signal was widely observed on all chromosomes (Fig. 8a, arrowhead 8). In round spermatids at stage I to elongated spermatids at stage X, strong dot-like signals were observed on the chromocenter in addition to weak and broad signals in the nucleus (Fig. 8a, arrowhead 12), while strong dot-like signals of H3K27me2 disappeared in elongated spermatids at stage X (Fig. 7a, arrowhead 12). Thus, localization of H3K27me3 modification seems to start from leptotene cells, the same timing as H3K27me2, and to accumulate on the chromocenter in early round spermatids (Fig. 8b).

**Fig. 8.**
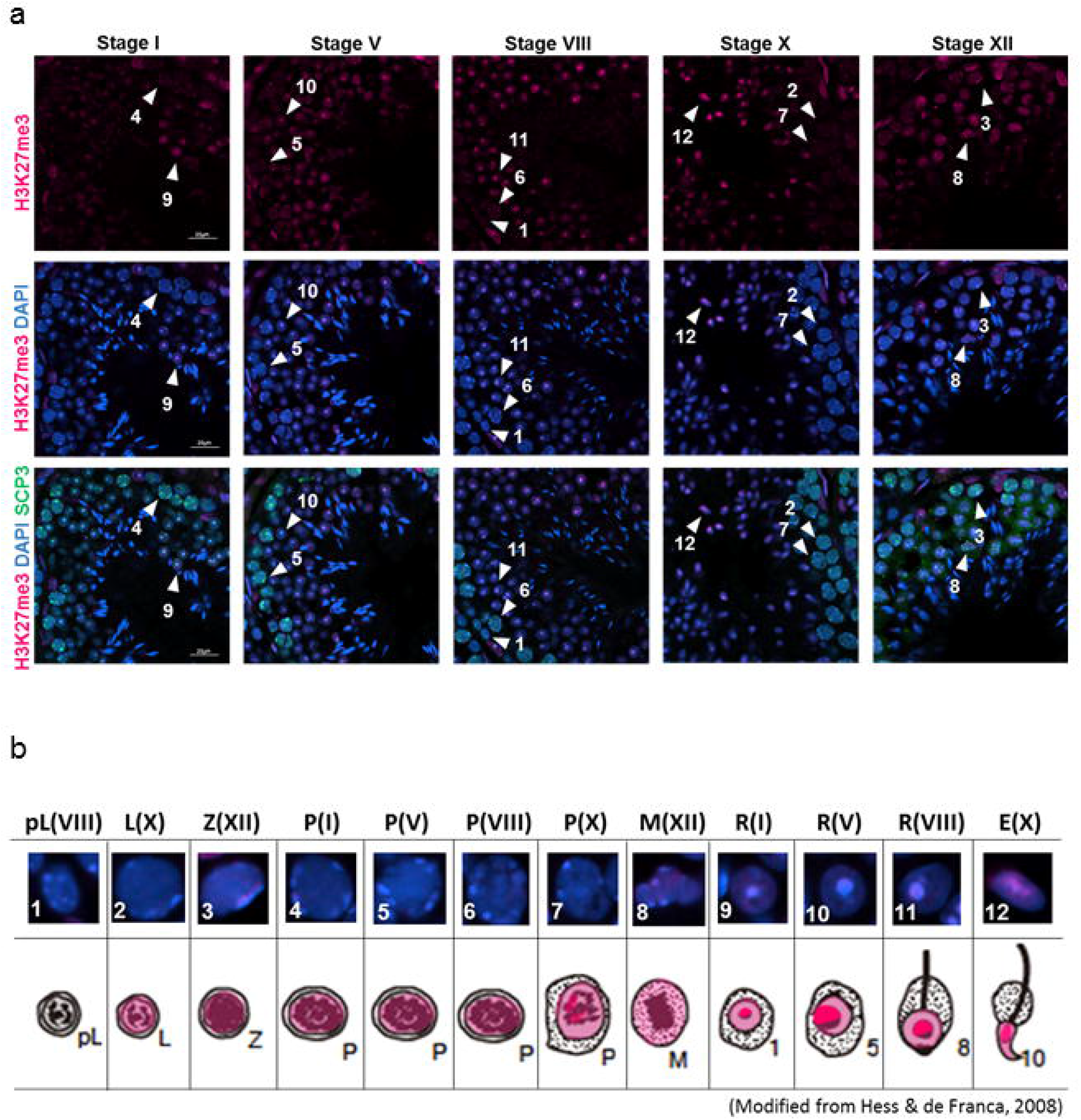
Localization of H3K27me3 in male germ line cells in the young (3 M) testis. (a) Representative confocal images of H3K27me3 (magenta) and SCP3 (green) in the seminiferous epithelium in stages I, V, VIII, X and XII. Nuclei were counterstained with DAPI (blue). The number with arrowhead means as the followings: 1; Preleptotene spermatocyte at stage VIII, 2; Leptotene spermatocyte at stage X, 3; Zygotene spermatocyte at stage XII, 4-7; Pachytene spermatocyte at stage I, V, VIII and X respectively, M; M phase cells at stage XII, 9-11; Round spermatid at stage I, V and VIII respectively, 12; Elongated spermatid at stage X. Scale bar: 20 μm. (b) Magnified images of the cells indicated by arrowheads in (a), and subcellular localization of H3K27me3 in the young testis is shown in magenta in a graphical summary. pL: Preleptotene spermatocyte, L: Leptotene spermatocyte, Z: Zygotene spermatocyte, P: Pachytene spermatocyte, M: M phase, R: Round spermatid, E: Elongated spermatid. Each parenthesis represents the stage of spermatogenesis.

The above localization patterns of histone modifications during spermatogenesis are shown in Table 1. In summary, H3K4me2 was detected in preleptotene cells to elongated spermatids, and H3K27ac was detected in leptotene cells to elongated spermatids. These two modifications were detected at the early stage of spermatogenesis, while other activation markers were detected at the later stage of spermatogenesis. All repressive makers examined in this study were detected at the early stage of spermatogenesis. All activation markers except H3K4me3 showed specific accumulation on XY body in pachytene cells, or a sex chromosome in round spermatids, while all repressive makers were accumulated on the chromocenter (i.e., constitutive heterochromatin). Only H3K9me3 was accumulated on both sex chromosomes and the chromocenter.

**Table 1.** Summarized expression levels of histone modifications during spermatogenesis and their alteration by aging. Expression levels of histone modifications that activate (red) and inhibit (blue) gene expression are shown. Each expression level of male germ line cells derived from young mice is indicated in + (faint), ++ (fair), +++ (bright), ++++ (very strong) or – (not detected). Arrows show expression level of male germ line cells derived from aged mice as compared to that of young mice. Single up/down arrows indicate that it was increased/decreased less than 20 % compared to young. Double up/down arrows indicate that it was increased/decreased more than 20 % compared to young. Right arrows indicate that there were no significant differences between young and aged (*p < 0.05, **p < 0.01). Specific localization or accumulation is described as specific notes.

|  |  | pL(VIII) | L(X) | Z(XII) | P(I) | P(V) | P(VIII) | P(X) | M(XII) | R(I) | R(V) | R(VIII) | E(X) | Specific note |
| --- | --- | --- | --- | --- | --- | --- | --- | --- | --- | --- | --- | --- | --- | --- |
| H3K4me2 | Young | +++ | +++ | ++ | ++ | ++ | + | ++ | ++ | ++ | +++ | ++ | ++ | Accumulation on XY body |
|  | Aged | ↓↓** | ↓↓** | ↓↓** | ↓↓** | ↓↓** | ↓↓** | ↓↓** | ↓↓** | ↓↓** | ↓↓** | ↓↓** | ↓↓** |  |
| H3K4me3 | Young | - | - | - | - | - | - | - | ++ | ++ | ++ | + | + |  |
|  | Aged | - | - | - | - | - | - | - | ↓↓** | → | ↓* | → | → |  |
| H3K27ac | Young | ++++ | + | + | +++ | ++ | ++ | +++ | + | +++ | +++ | + | ++++ | Accumulation on XY body and sex chromosomes |
|  | Aged | → | ↓* | → | → | ↓* | → | ↓** | → | → | → | → | ↑* |  |
| H3K79me2 | Young | - | - | - | - | - | + | + | ++ | ++ | +++ | ++++ | +++ | Accumulation on sex chromosomes |
|  | Aged | - | - | - | - | - | → | ↑** | → | → | → | ↑** | ↑↑** |  |
| H3K79me3 | Young | - | - | - | - | - | - | - | +++ | - | + | ++ | +++ | Accumulation on sex chromosomes |
|  | Aged | - | - | - | - | - | - | - | ↑↑** | - | → | ↓* | ↓* |  |
| H3K9me3 | Young | ++ | ++ | ++ | ++ | ++ | ++ | ++ | ++ | ++ | ++ | +++ | +++ | Accumulation on sex chromosomes and heterochromatin |
|  | Aged | ↓↓** | ↓↓** | ↓↓** | ↓↓** | ↓↓** | ↓↓** | ↓↓** | → | ↓** | ↑* | ↓** | → |  |
| H3K27me2 | Young | - | ++ | + | + | ++ | +++ | +++ | ++ | + | ++ | + | +++ | Accumulation on heterochromatin |
|  | Aged | - | ↑↑** | ↑↑** | ↑↑** | ↑↑** | ↓** | ↑↑** | ↑↑** | ↑↑** | ↑↑** | ↓** | ↑↑** |  |
| H3K27me3 | Young | - | + | + | + | + | + | + | ++ | ++ | + | + | ++ | Accumulation on heterochromatin |
|  | Aged | - | ↓* | ↑↑** | ↑↑** | ↑↑** | ↑↑** | ↓** | → | ↑↑** | ↑↑** | → | → |  |

### 2. Semi-quantitative analyses of histone modifications in young and aged testes

We next performed semi-quantified analyses to examine whether intensity of histone modifications are changed in young and aged testes. The signal of histone modification was measured by ImageJ. The intensity of histone modification was normalized by that of DAPI in the nucleus or chromosomes.

#### (1) Histone modifications that activate gene expression

No difference was observed in localization patterns of H3K4me2/3, H3K27ac, H3K79me2/3, H3K9me3 and H3K27me2/3 in young and aged testes (Fig. 9a, 10a, 11a, 12a, 13a, 14a, 15a and 16a). However, semi-quantitative analyses revealed difference in intensity levels among histone modifications observed.

**Fig. 9.**
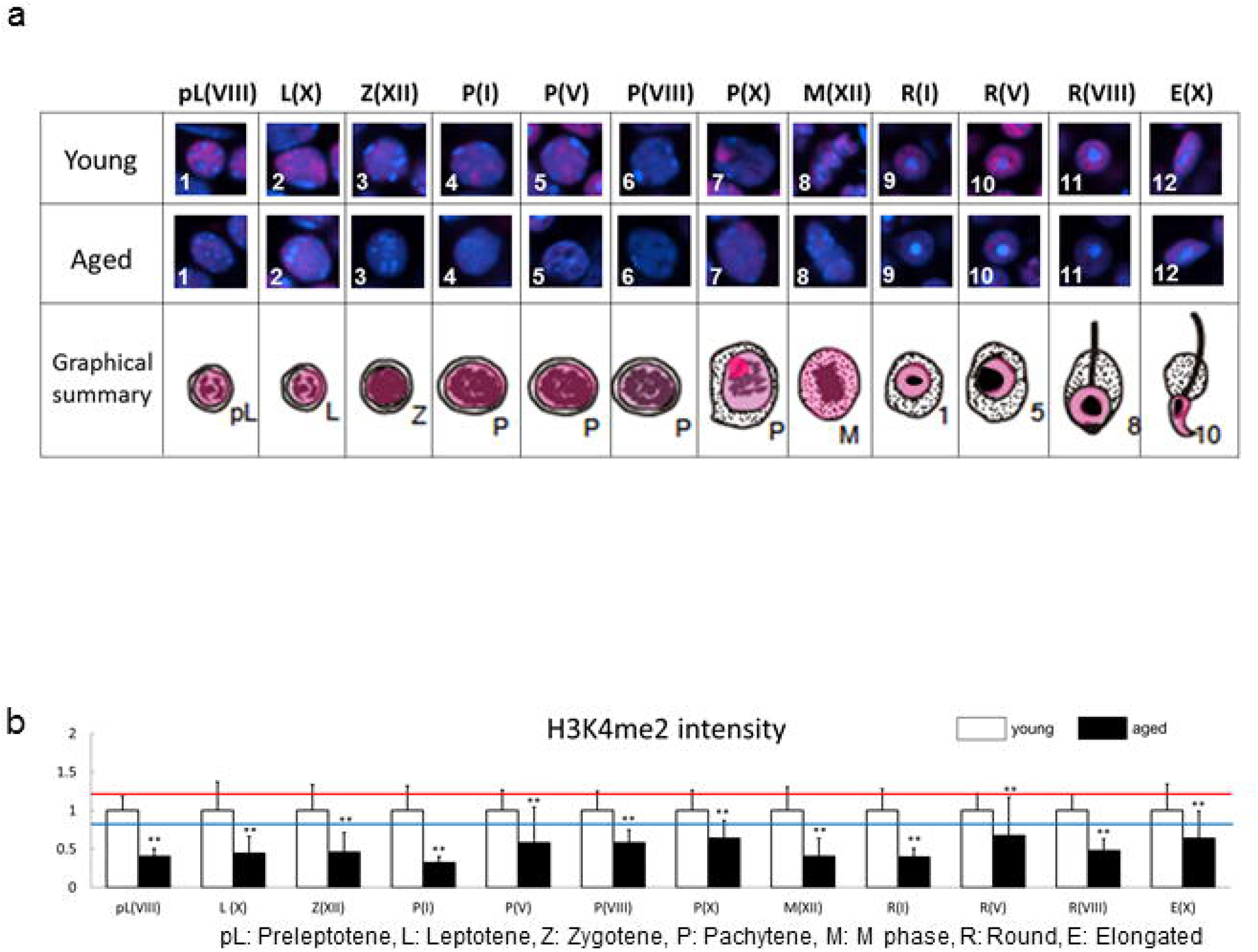
Comparison of localization patterns and intensity of H3K4me2 in male germ line cells in young (3M) and aged (12M) testes. (a) Representative confocal images of H3K4me2 (magenta) in male germ line cells in young and aged testes. Nuclei were counterstained with DAPI (blue). Summarized illustration showing subcellular localization of H3K4me2 (magenta) in male germ line cells in young and aged testes. The number means as the followings: 1; Preleptotene spermatocyte at stage VIII, 2; Leptotene spermatocyte at stage X, 3; Zygotene spermatocyte at stage XII, 4-7; Pachytene spermatocyte at stage I, V, VIII and X respectively, M; M phase cells at stage XII, 9-11; Round spermatid at stage I, V and VIII respectively, 12; Elongated spermatid at stage X. (b) Semi-quantitative analysis of H3K4me2 in male germ line cells in young and aged testes. Black bars show relative intensity of aged cells against young ones (**p < 0.01, *p < 0.05, n.d., not detected). The red and blue lines mean 20% increased and decrease to the young testis, respectively. pL: Preleptotene spermatocyte, L: Leptotene spermatocyte, Z: Zygotene spermatocyte, P: Pachytene spermatocyte, M: M phase, R: Round spermatid, E: Elongated spermatid. Each parenthesis represents the stage of spermatogenesis.

##### (i) H3K4me2

Intensity levels were lower in all the cells of the aged testis than in the young testis (Fig. 9b).

##### (ii) H3K4me3

Intensity levels were lower in M-phase spermatocytes at stage XII and round spermatids at stage V of the aged testis than in the young testis (Fig. 10b). In other cells, there were no statistical differences.

**Fig. 10.**
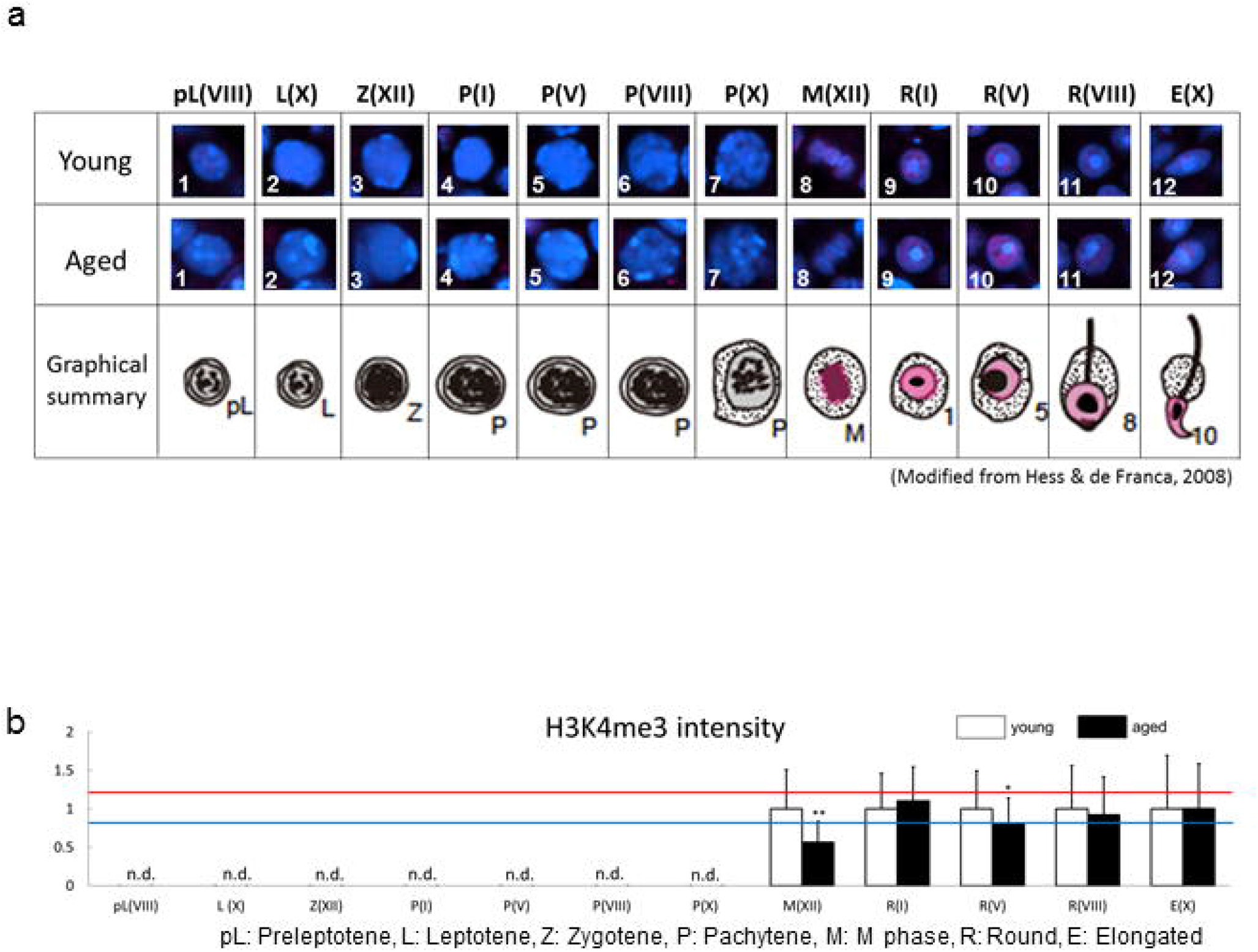
Comparison of localization patterns and intensity of H3K4me3 in male germ line cells in young (3M) and aged (12M) testes. (a) Representative confocal images of H3K4me3 (magenta) in male germ line cells in young and aged testes. Nuclei were counterstained with DAPI (blue). Summarized illustration showing subcellular localization of H3K4me3 (magenta) in male germ line cells in young and aged testes. The number means as the followings: 1; Preleptotene spermatocyte at stage VIII, 2; Leptotene spermatocyte at stage X, 3; Zygotene spermatocyte at stage XII, 4-7; Pachytene spermatocyte at stage I, V, VIII and X respectively, M; M phase cells at stage XII, 9-11; Round spermatid at stage I, V and VIII respectively, 12; Elongated spermatid at stage X. (b) Semi-quantitative analysis of H3K4me3 in male germ line cells in young and aged testes. Black bars show relative intensity of aged cells against young ones (**p < 0.01, *p < 0.05, n.d., not detected). The red and blue lines mean 20% increased and decrease to the young testis, respectively. pL: Preleptotene spermatocyte, L: Leptotene spermatocyte, Z: Zygotene spermatocyte, P: Pachytene spermatocyte, M: M phase, R: Round spermatid, E: Elongated spermatid. Each parenthesis represents the stage of spermatogenesis.

##### (iii) H3K27ac

Intensity levels were lower in leptotene cells at stage X, pachytene cells at stage V and X of the aged testis than in the young testis (Fig. 11b). However, higher intensity was detected in stage X elongated spermatids in the aged testis. In other cells, there were no statistical differences.

**Fig. 11.**
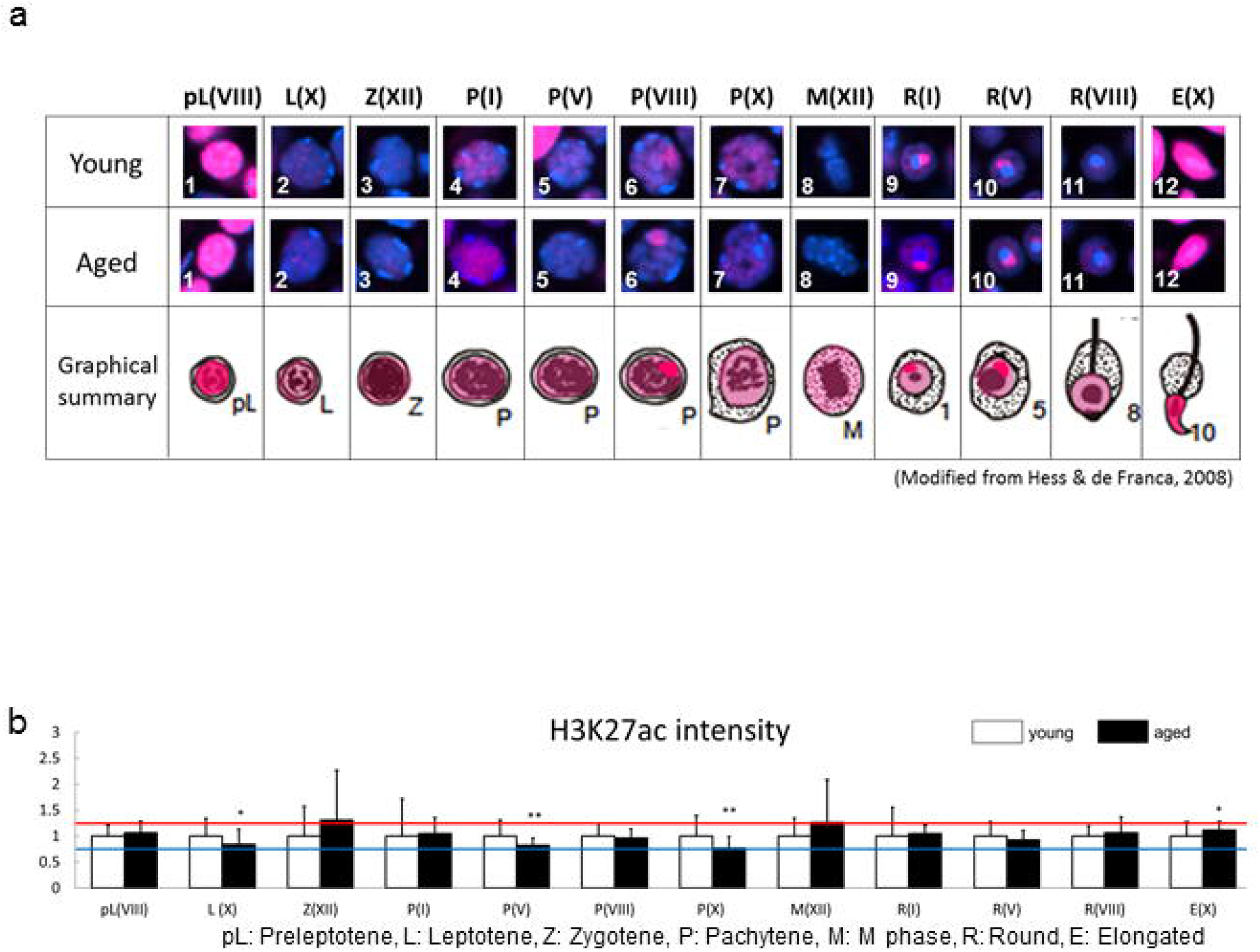
Comparison of localization patterns and intensity of H3K27ac in male germ line cells in young (3M) and aged (12M) testes. (a) Representative confocal images of H3K27ac (magenta) in male germ line cells in young and aged testes. Nuclei were counterstained with DAPI (blue). Summarized illustration showing subcellular localization of H3K27ac (magenta) in male germ line cells in young and aged testes. The number means as the followings: 1; Preleptotene spermatocyte at stage VIII, 2; Leptotene spermatocyte at stage X, 3; Zygotene spermatocyte at stage XII, 4-7; Pachytene spermatocyte at stage I, V, VIII and X respectively, M; M phase cells at stage XII, 9-11; Round spermatid at stage I, V and VIII respectively, 12; Elongated spermatid at stage X. (b) Semi-quantitative analysis of H3K27ac in male germ line cells in young and aged testes. Black bars show relative intensity of aged cells against young ones (**p < 0.01, *p < 0.05, n.d., not detected). The red and blue lines mean 20% increased and decrease to the young testis, respectively. pL: Preleptotene spermatocyte, L: Leptotene spermatocyte, Z: Zygotene spermatocyte, P: Pachytene spermatocyte, M: M phase, R: Round spermatid, E: Elongated spermatid. Each parenthesis represents the stage of spermatogenesis.

##### (iv) H3K79me2

Comparing with the young testis, higher intensity was detected in pachytene cells at stage X, round spermatids at stage VIII, and elongated spermatids at stage X (Fig. 12b) of the aged testis. However, other cells showed no difference in the intensity level.

**Fig. 12.**
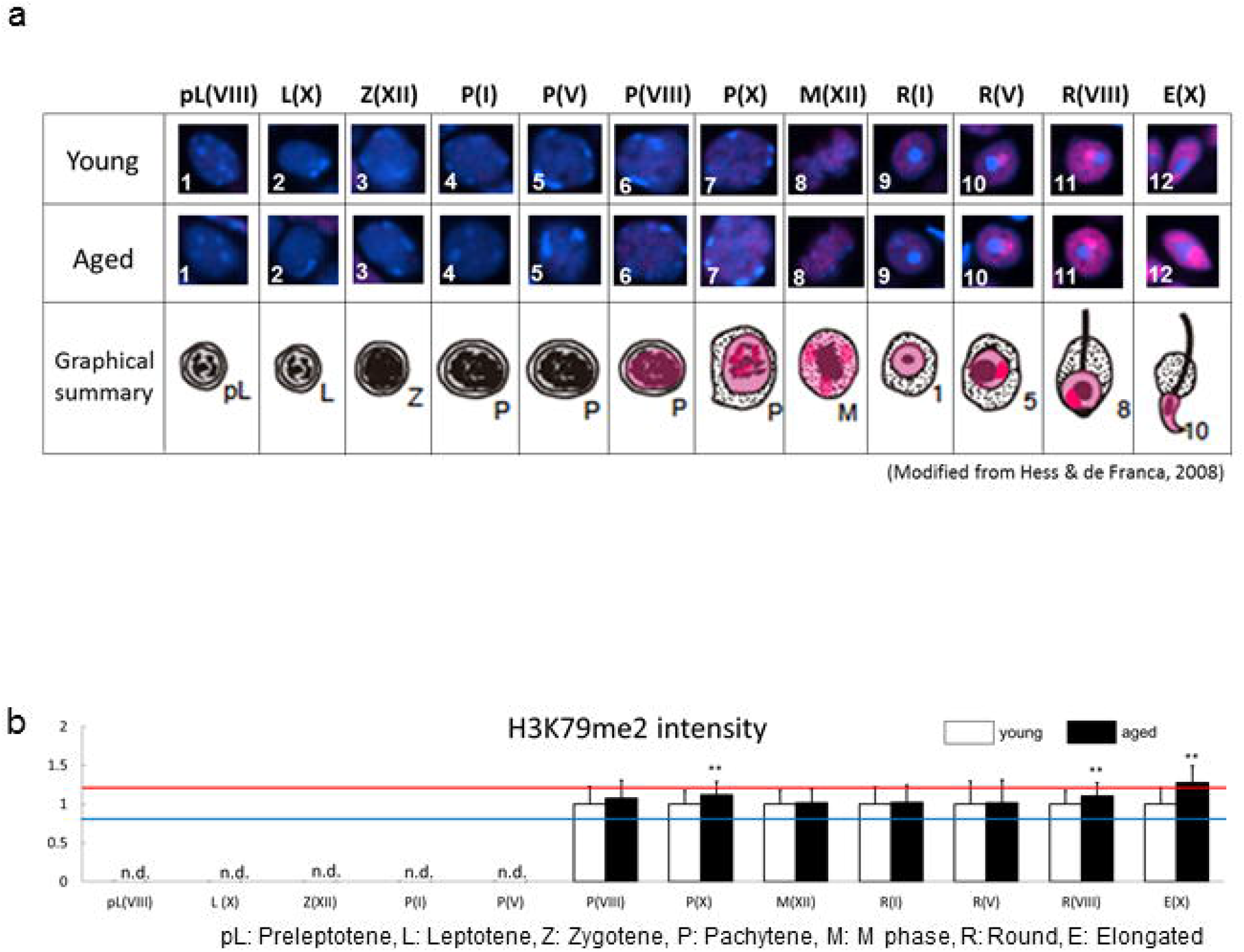
Comparison of localization patterns and intensity of H3K79me2 in male germ line cells in young (3M) and aged (12M) testes. (a) Representative confocal images of H3K79me2 (magenta) in male germ line cells in young and aged testes. Nuclei were counterstained with DAPI (blue). Summarized illustration showing subcellular localization of H3K79me2 (magenta) in male germ line cells in young and aged testes. The number means as the followings: 1; Preleptotene spermatocyte at stage VIII, 2; Leptotene spermatocyte at stage X, 3; Zygotene spermatocyte at stage XII, 4-7; Pachytene spermatocyte at stage I, V, VIII and X respectively, M; M phase cells at stage XII, 9-11; Round spermatid at stage I, V and VIII respectively, 12; Elongated spermatid at stage X. (b) Semi-quantitative analysis of H3K79me2 in male germ line cells in young and aged testes. Black bars show relative intensity of aged cells against young ones (**p < 0.01, *p < 0.05, n.d., not detected). The red and blue lines mean 20% increased and decrease to the young testis, respectively. pL: Preleptotene spermatocyte, L: Leptotene spermatocyte, Z: Zygotene spermatocyte, P: Pachytene spermatocyte, M: M phase, R: Round spermatid, E: Elongated spermatid. Each parenthesis represents the stage of spermatogenesis.

##### (v) H3K79me3

Higher intensity was detected in M-phase spermatocytes at stage XII of the aged testis than in the young testis, while lower intensity was observed in round spermatids at stage VIII and elongate spermatids at stage X (Fig. 13b).

**Fig. 13.**
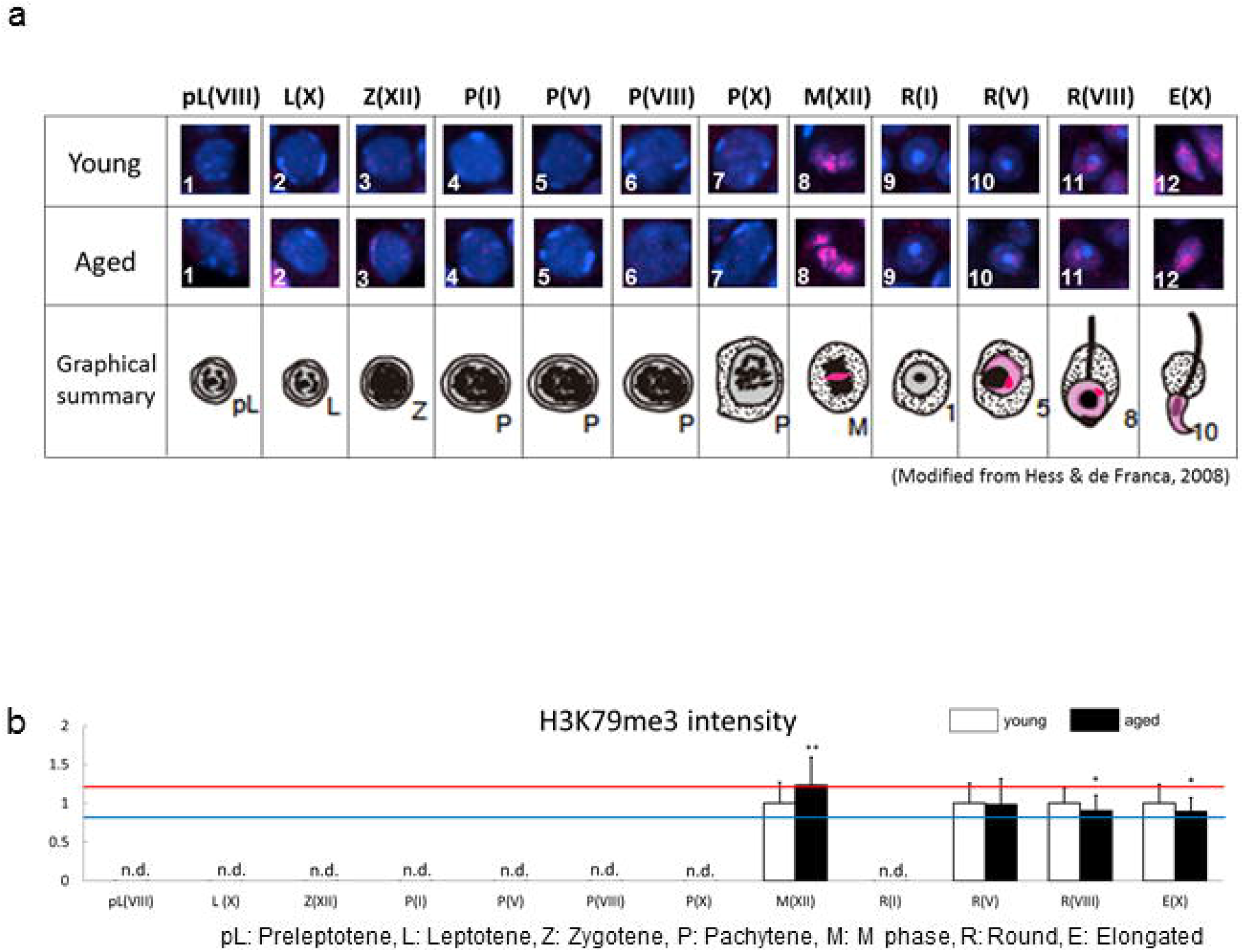
Comparison of localization patterns and intensity of H3K79me3 in male germ line cells in young (3M) and aged (12M) testes. (a) Representative confocal images of H3K79me3 (magenta) in male germ line cells in young and aged testes. Nuclei were counterstained with DAPI (blue). Summarized illustration showing subcellular localization of H3K79me3 (magenta) in male germ line cells in young and aged testes. The number means as the followings: 1; Preleptotene spermatocyte at stage VIII, 2; Leptotene spermatocyte at stage X, 3; Zygotene spermatocyte at stage XII, 4-7; Pachytene spermatocyte at stage I, V, VIII and X respectively, M; M phase cells at stage XII, 9-11; Round spermatid at stage I, V and VIII respectively, 12; Elongated spermatid at stage X. (b) Semi-quantitative analysis of H3K79me3 in male germ line cells in young and aged testes. Black bars show relative intensity of aged cells against young ones (**p < 0.01, *p < 0.05, n.d., not detected). The red and blue lines mean 20% increased and decrease to the young testis, respectively pL: Preleptotene spermatocyte, L: Leptotene spermatocyte, Z: Zygotene spermatocyte, P: Pachytene spermatocyte, M: M phase, R: Round spermatid, E: Elongated spermatid. Each parenthesis represents the stage of spermatogenesis.

#### (2) Histone modifications that inhibit gene expression

##### (i) H3K9me3

Intensity levels were lower in almost all the cells of the aged testis than in the young testis (Fig. 14b). However, higher intensity was detected in stage V round spermatids in the aged testis. There were no statistical differences in other cell types.

**Fig. 14.**
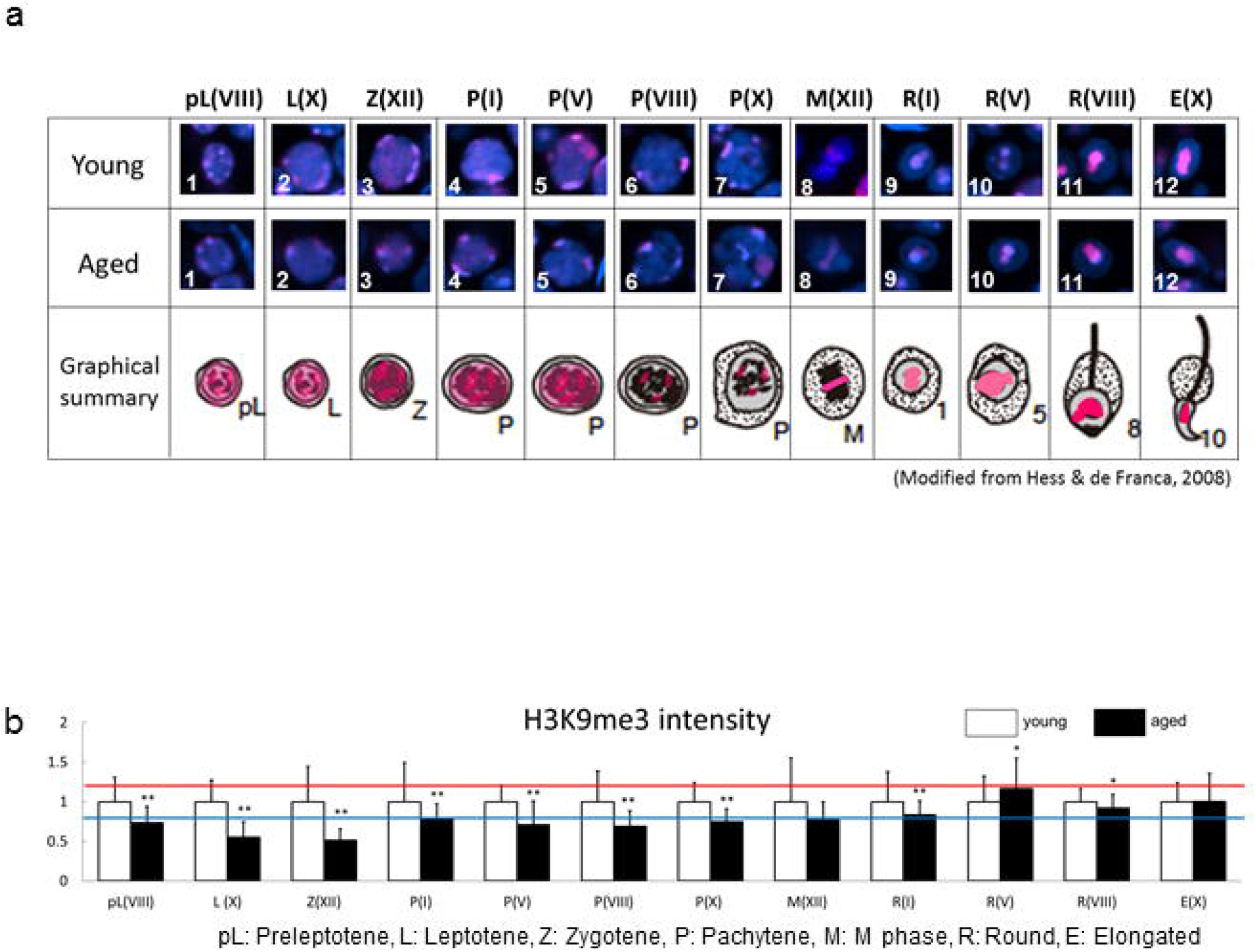
Comparison of localization patterns and intensity of H3K9me3 in male germ line cells in young (3M) and aged (12M) testes. (a) Representative confocal images of H3K9me3 (magenta) in male germ line cells in young and aged testes. Nuclei were counterstained with DAPI (blue). Summarized illustration showing subcellular localization of H3K9me3 (magenta) in male germ line cells in young and aged testes. The number means as the followings: 1; Preleptotene spermatocyte at stage VIII, 2; Leptotene spermatocyte at stage X, 3; Zygotene spermatocyte at stage XII, 4-7; Pachytene spermatocyte at stage I, V, VIII and X respectively, M; M phase cells at stage XII, 9-11; Round spermatid at stage I, V and VIII respectively, 12; Elongated spermatid at stage X. (b) Semi-quantitative analysis of H3K9me3 in male germ line cells in young and aged testes. Black bars show relative intensity of aged cells against young ones (**p < 0.01, *p < 0.05, n.d., not detected). The red and blue lines mean 20% increased and decrease to the young testis, respectively pL: Preleptotene spermatocyte, L: Leptotene spermatocyte, Z: Zygotene spermatocyte, P: Pachytene spermatocyte, M: M phase, R: Round spermatid, E: Elongated spermatid. Each parenthesis represents the stage of spermatogenesis.

##### (ii) H3K27me2

Intensity levels were lower in pachytene cells at stage VIII and round spermatids at stage VIII of the aged testis than in the young testis (Fig. 15b). However, other cells showed higher intensity in the aged testis.

**Fig. 15.**
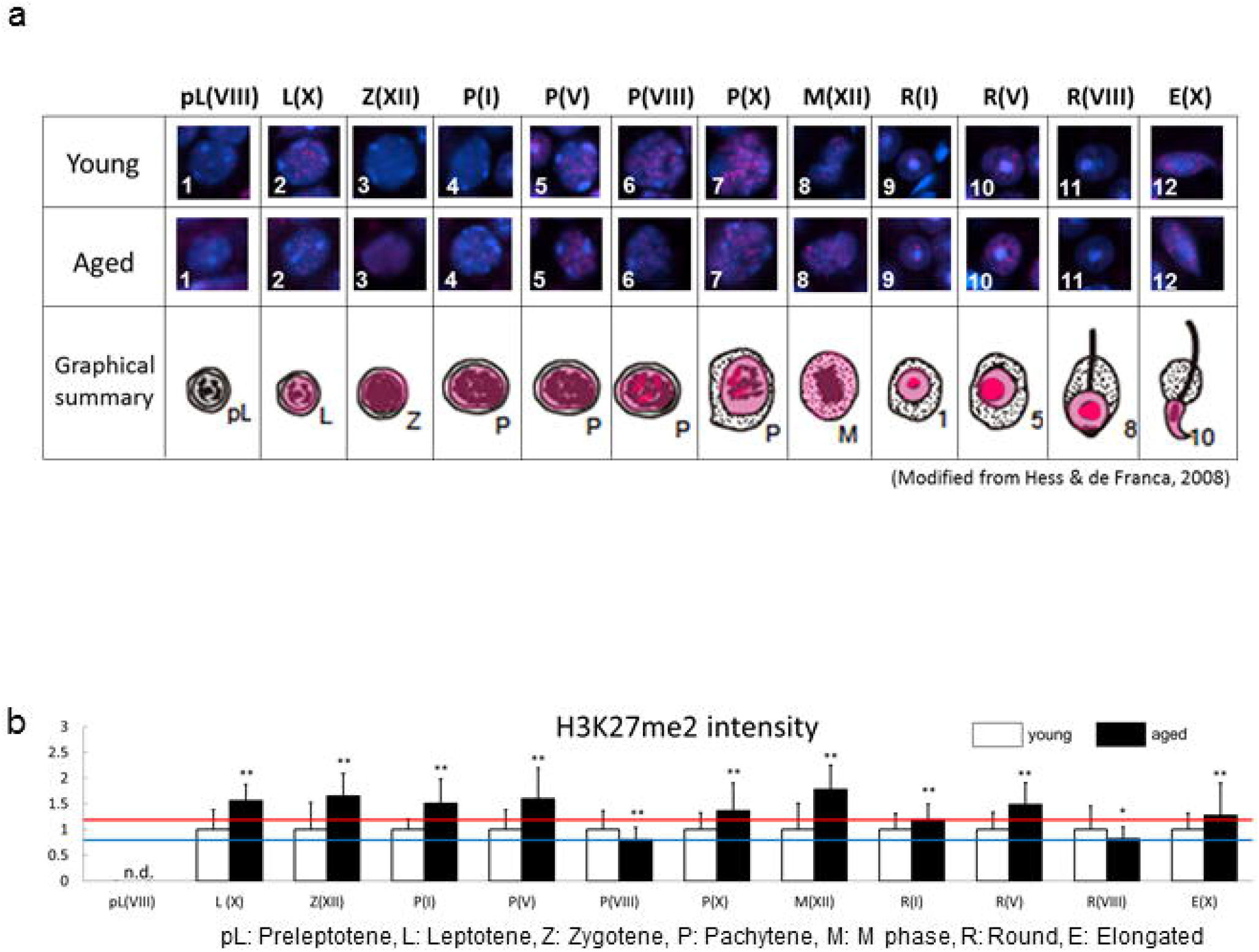
Comparison of localization patterns and intensity of H3K27me2 in male germ line cells in young (3M) and aged (12M) testes. (a) Representative confocal images of H3K27me2 (magenta) in male germ line cells in young and aged testes. Nuclei were counterstained with DAPI (blue). Summarized illustration showing subcellular localization of H3K27me2 (magenta) in male germ line cells in young and aged testes. The number means as the followings: 1; Preleptotene spermatocyte at stage VIII, 2; Leptotene spermatocyte at stage X, 3; Zygotene spermatocyte at stage XII, 4-7; Pachytene spermatocyte at stage I, V, VIII and X respectively, M; M phase cells at stage XII, 9-11; Round spermatid at stage I, V and VIII respectively, 12; Elongated spermatid at stage X. (b) Semi-quantitative analysis of H3K27me2 in male germ line cells in young and aged testes. Black bars show relative intensity of aged cells against young ones (**p < 0.01, *p < 0.05, n.d., not detected). The red and blue lines mean 20% increased and decrease to the young testis, respectively. pL: Preleptotene spermatocyte, L: Leptotene spermatocyte, Z: Zygotene spermatocyte, P: Pachytene spermatocyte, M: M phase, R: Round spermatid, E: Elongated spermatid. Each parenthesis represents the stage of spermatogenesis.

##### (iii) H3K27me3

Similar to the result of H3K27me2, almost stages of cells showed higher intensity in the aged testis (Fig. 16b). In stage X leptotene cells and stage VIII pachytene cells, lower intensity was detected in the aged testis. There were no statistical differences in other cell types.

**Fig. 16.**
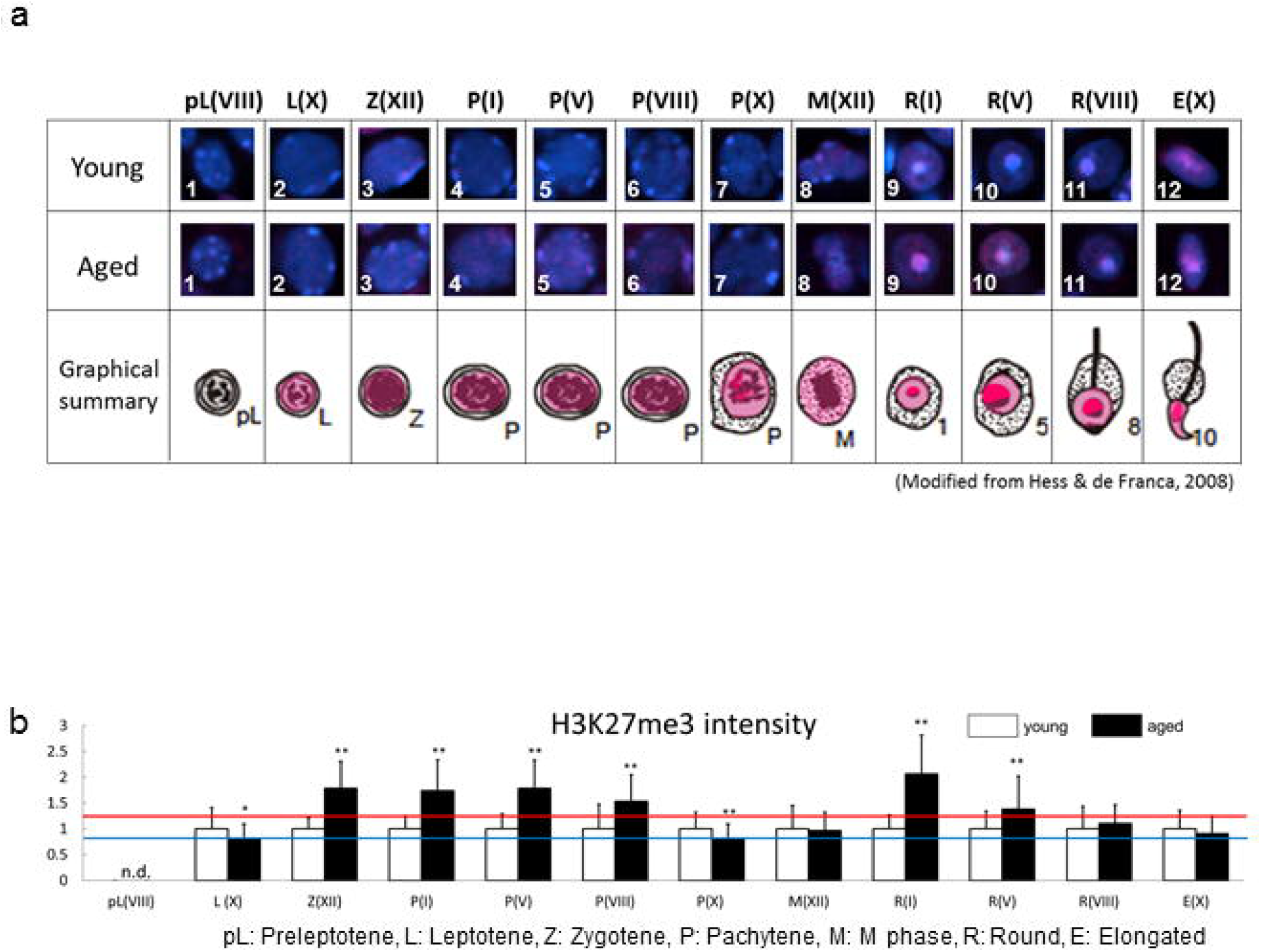
Comparison of localization patterns and intensity of H3K27me3 in male germ line cells in young (3M) and aged (12M) testes. (a) Representative confocal images of H3K27me3 (magenta) in male germ line cells in young and aged testes. Nuclei were counterstained with DAPI (blue). Summarized illustration showing subcellular localization of H3K27me3 (magenta) in male germ line cells in young and aged testes. The number means as the followings: 1; Preleptotene spermatocyte at stage VIII, 2; Leptotene spermatocyte at stage X, 3; Zygotene spermatocyte at stage XII, 4-7; Pachytene spermatocyte at stage I, V, VIII and X respectively, M; M phase cells at stage XII, 9-11; Round spermatid at stage I, V and VIII respectively, 12; Elongated spermatid at stage X. (b) Semi-quantitative analysis of H3K27me3 in male germ line cells in young and aged testes. Black bars show relative intensity of aged cells against young ones (**p < 0.01, *p < 0.05, n.d., not detected). The red and blue lines mean 20% increased and decrease to the young testis, respectively. pL: Preleptotene spermatocyte, L: Leptotene spermatocyte, Z: Zygotene spermatocyte, P: Pachytene spermatocyte, M: M phase, R: Round spermatid, E: Elongated spermatid. Each parenthesis represents the stage of spermatogenesis.

The above intensity of histone modifications in the young and aged testis is shown in Table 1. In summary, histone modifications were altered by aging and the alteration patterns were different in individual histone modifications. Among the repressive marks, H3K9me3 level was decreased in the aged testis, while H3K27me2 and H3K27me3 level was increased during spermatogenesis. On the other hand, the intensity level of an activation mark, H3K4me2, was drastically decreased in the aged testis during spermatogenesis. Other histone modifications also showed differences at some stages.

### 3. Accumulation of H3K79me3 on sex chromosomes

As described above, H3K79me3 was detected in M-phase spermatocytes, round spermatids and elongated spermatids. It is previously reported that H3K79me3 accumulates on XY body in late pachytene cells and on a sex chromosome in round spermatids (13). In this study, H3K79me3 signal was strongly detected on specific chromosomes in M-phase spermatocytes at stage XII (Fig. 17a). Therefore, we suspected that the chromosome on which H3K79me3 accumulated might be sex chromosomes.

**Fig. 17.**
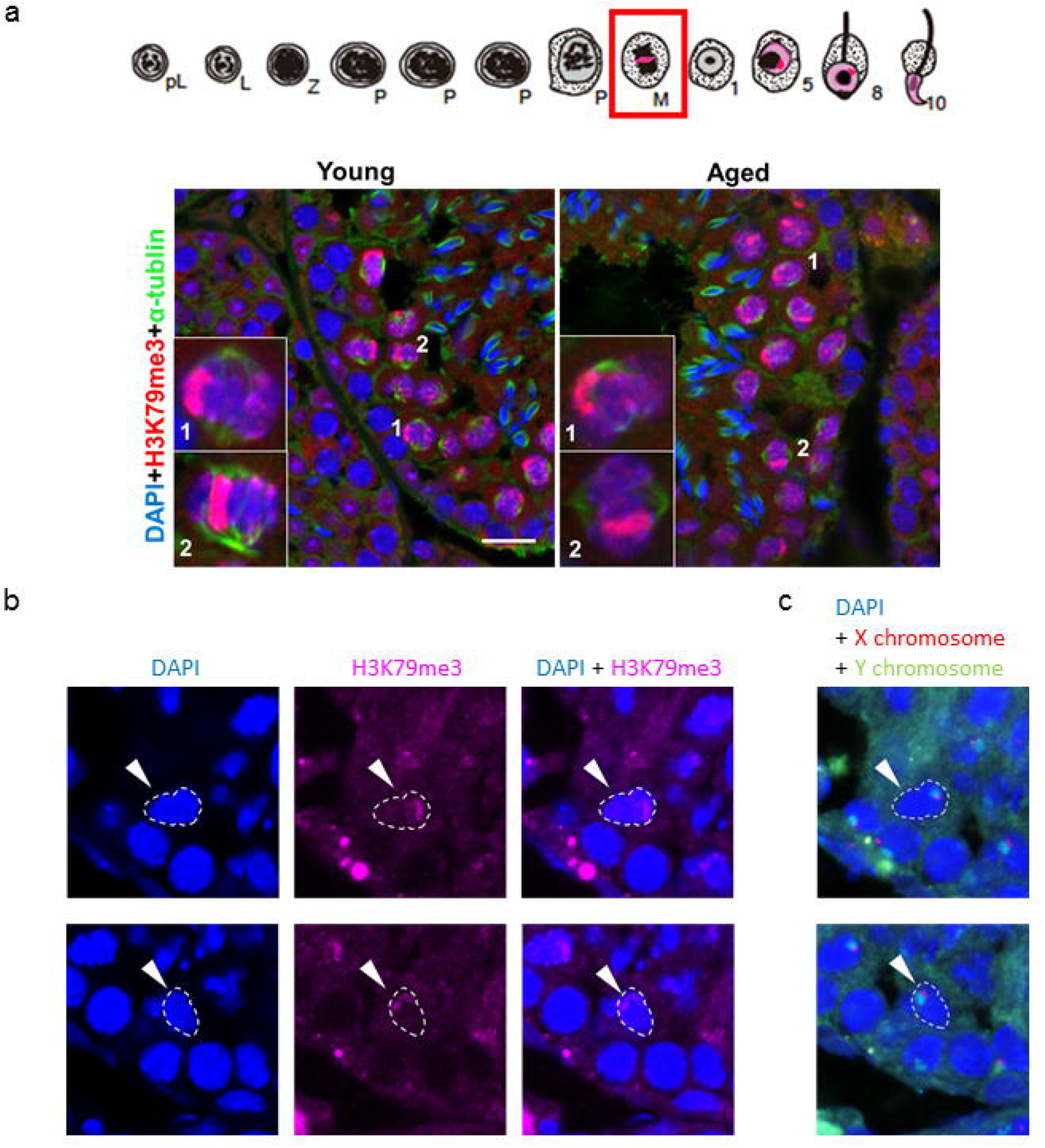
H3K79me3 localization on sex chromosomes in the spermatocyte at M phase. (a) Summarized illustration showing subcellular localization of H3K79me3 (magenta). Lower panels show representative confocal images of H3K79me3 (magenta) and tubulin (green) in M phase cells (red square in the summarized illustration) in young and aged testes. Nuclei were counterstained with DAPI (blue). Magnified images of the cells indicated with numbers are shown in the box. (b) Representative confocal images of H3K79me3 (magenta) in M phase cells. Nuclei were counterstained with DAPI (blue). M phase cells are indicated with dotted line and arrowheads. (c) Representative images of FISH showing X (red) and Y (green) chromosomes in M phase cells on the same section as (b). Nuclei were counterstained with Hoechst (blue). The same M phase cells as (b) are indicated with dotted lines and arrowheads.

To identify the sex chromosome, we established a new protocol by combining immunostaining together with fluorescence in situ hybridization (FISH) on the same slide. First, immunostaining was performed to detect H3K79me3 signal in M-phase spermatocytes (Fig. 17b). After images were captured using confocal microscopy, FISH was performed using probes that recognize X and Y chromosomes, respectively, followed by capturing FISH images of the same slides to obtain triple signals (Fig. 17c). Signals of H3K79me3 was overlapped with those of X and Y chromosomes at the same position within the nucleus of M-phase spermatocytes. Finally, we revealed that H3K79me3 was specifically accumulated on the sex chromosomes in M-phase spermatocytes. This result is consistent with another study that suggest the accumulation of H3K79me3 on the sex chromosomes in late pachytene cells and round spermatids (13).

## Discussion

In this study, we have made a catalog showing transition of histone modification localization during spermatogenesis and its alteration caused by aging. Individual histone modifications showed distinct localization patterns and different timing to start their expression. Even among the histone modifications that have similar functions showed different localization patterns. Moreover, intensity of histone modifications was altered by aging. Aged testes showed differential increased/decreased signals of histone modifications at different stages. Therefore, responses to aging seem to be different among histone modifications. We also identified a chromosome in which H3K79me3 accumulated in the M-phase spermatocyte. H3K79me3 was accumulated on sex chromosomes in M-phase spermatocytes and round spermatids. Morphologically, it is revealed that not only H3K79me3, but also other histone modifications such as H3K79me2, H3K27ac and H3K9me3 accumulated on putative sex chromosomes or XY body. The intensity of these histone modification specifically accumulated on the putative sex chromosomes or XY body was also altered by aging at some stages. Therefore, epigenetic regulation of histone modifications may be different between sex chromosomes and other chromosomes, which may also be affected by aging.

In the present study, histone modifications that inhibit gene expression showed more drastic changes in the aged testis. At the pachytene stage of meiosis I, genes on sex chromosomes are transcriptionally shut down by meiotic sex chromosome inactivation (MSCI), and reactivated after meiosis (14). It is reported that H3K9me3 is involved in silencing of sex chromosomes in meiosis and that if there is no enrichment of H3K9me3, remodeling and silencing of sex chromosomes failed and cause germ cell apoptosis (12). In post-meiotic spermatids, 87% of X-linked genes remain suppressed, while autosomes are largely active (15). Moreover, spermatids form a distinct post-meiotic sex chromatin (PMSC) area that is enriched with H3K9me3 (15, 16). Thus, H3K9me3, a repressive mark, has various roles during meiosis and in the post-meiosis phase. In our results, H3K9me3 appeared from preleptotene cells to elongated spermatids, and drastically decreased in the aged testis during meiosis and in most of the stages of round spermatids. It is assumed in the aged testis that failure of correct chromosome segregation during meiosis or leaky expression of X-linked genes might occur in the post-meiotic spermatids even though the male germ line cells safely pass meiosis.

Another repressive mark, H3K27me3, appeared from leptotene cells to elongated spermatids. A study has reported that H3K27ac may act as an antagonist of Polycomb-mediated silencing through inhibition of H3K27me3 (17). H3K27ac showed signals in all stages of the male germ line cells examined, especially in preleptotene cells and elongated spermatids. It is very interesting that H3K27ac and H3K27me3 showed exclusive localization in round spermatids; the former was accumulated on the sex chromosome, while H3K27me3 on the chromocenter, a heterochromatin region. Therefore, the absence of H3K27ac (an activation mark) and presence of H3K27me3 (a repressive mark) in the chromocenter may be reasonable in regard with gene regulation. We found that H3K27me3 was dramatically increased in the aged testis, while H3K27ac showed slightly decreased at certain stages. H3K27me3 inhibits transcription by inducing a compact chromatin structure (18, 19). Thus, gene expression influenced by H3K27me3 might be more repressed in the aged testis.

It is not easy to elucidate the function of histone modification and its alteration by aging. However, progenia models can be used to know aging-induced phenotypes and their underlying mechanisms. Hutchinson-Gilford progeria syndrome (HGPS) patients show premature aging, and cultured cells derived from HGPS patients exhibit an increase of H4K20me3 and a reduction of both H3K9me3 and H3K27me3 (20), i.e., histone modifications inducing heterochromatin regions (21). Although this result seems to be rather puzzling based on our general knowledge about parallel regulation of H3K9me3 and H4K20me3, it might be considered that heterochromatin regions are decreased in HGPS patients. Indeed, loss of heterochromatin is considered to be one of the mechanisms involved in aging (22, 23). However, H3K27me2/3 was increased at the most of stages in the aged testis. A similar increase of H3K27me3 due to aging is observed in quiescent muscle stem cells (24). Therefore, it may reasonable to assume that increase/decrease of H3K27me3 may depend on tissues or cell types. One possible reason of these difference can be explained by activities of histone methyltransferases or demethylases, some of which have tissue- or cell line-specific function (25, 26). Polycomb-repressive complex 2 (PRC2) that catalyzes the methylation of H3K27 transcriptionally represses soma-specific genes and facilitates homeostasis and differentiation during mammalian spermatogenesis (27). This specificity can be associated with different alteration patterns of histone modification in each tissue or cell line. It might be interesting to know whether H3K27me3 is increased by PRC2 and affected to spermatogenesis.

The sex chromosomes seemed to be unique in localization of histone modifications. In this study, we confirmed accumulation of H3K79me3 on the sex chromosomes in spermatocytes at M phase. A similar localization of H3K9me3 has been shown in a previous study (12). Moreover, H3K27ac was accumulated on XY body and sex chromosomes in spermatocytes and in round spermatids, respectively. H3K79me2 also showed a signal on the sex chromosomes in round spermatids. We found that intensity of these histone modifications specifically accumulated on the sex chromosomes and/or XY body were also affected by aging at certain stages. What do those results mean?

According to the database collecting risk genes for autism (SFARI at June 25, 2019), X chromosome has 81 risk genes, which is comparable to 95 and 89 risk genes on the larger chromosomes 1 and 2, respectively. It may be reasonable to assume that these three chromosomes have a large number of autism risk genes depending on their size. However, X chromosome has 16 “syndromic” genes, whereas chromosome 1 and 2 have only four and five, respectively. Enrichment of the autism risk genes, especially syndromic ones, on X chromosome might explain severer symptoms in female patients because X chromosome of father inevitably transmitted only to a daughter. Therefore, more detailed analyses would be interesting to focus on target genes of histone modifications that located on X chromosome.

Our results demonstrated aging-related alteration of histone modifications in spermatogenesis. In literatures, the possibility for the effect of paternal epigenetic factors on offspring’s health is suggested (28–31). Currently, however, there is no report explaining how the paternal epigenome is actually inherited to offspring. This will be a next important question in regard with the paternal impact on physical conditions of offspring. Although most of histone protein is replaced with protamine during spermatogenesis, some histone modifications in sperm are remained at promoter regions that are known to regulate embryonic development (6). Further studies are needed to reveal paternal aging impact on epigenetic mechanisms that may cause transgenerational influence on health and disease of progeny.

## Acknowledgements

The authors thank Prof. Yasuhisa Matsui for providing insightful comments and assistance, and Ms. Sayaka Makino for animal care. We are also grateful to all members of our laboratory for fruitful discussions and advice. This work was supported by KAKENHI (17K19372).

